# Biophysical characterization of Eag chaperones suggests the mechanism of effector transmembrane domain release

**DOI:** 10.1101/2025.03.20.644388

**Authors:** Matthew Van Schepdael, Iman Asakereh, Jake Colautti, Andrew J. Gierys, Shehryar Ahmad, Mazdak Khajehpour, John C. Whitney, Gerd Prehna

**Affiliations:** Department of Microbiology, University of Manitoba, Winnipeg Canada; Department of Chemistry, University of Manitoba, Winnipeg Canada; Department of Biochemistry and Biomedical Sciences, McMaster University, Hamilton, Canada

**Keywords:** Type VI secretion system, Eag, chaperone, membrane protein, stopped-flow, protein folding, X-ray crystallography

## Abstract

The type VI secretion system (T6SS) is a dynamic protein nanomachine found in Gram-negative bacteria that secretes toxic effectors into prey-cells. For secretion, effectors require chaperones or adaptors for proper loading onto the T6SS. Effector associated genes (Eags) are a family of T6SS chaperones that stabilize N-terminal transmembrane domains (TMDs) found in thousands of effectors. Eags are essential for secretion and inhibit effector TMDs from prematurely adopting a membrane-penetrative conformation. However, the mechanism of TMD release from its cognate Eag chaperone is unknown. Here, we take a biochemical and biophysical approach to probe the mechanism of TMD binding and dissociation from Eag chaperones. Using steady-state fluorescence, stopped-flow measurements, and bacterial competition assays, we compare the thermodynamics, kinetics, and *in vivo* chaperone function of wild-type and point-variant Eag-TMD complexes. Additionally, we solve an X-ray crystal structure of an Eag-TMD point-variant complex that captures an intermediate state of TMD release. Our data reveals the molecular features and specific residue contacts necessary for TMD binding and demonstrates the Eag conformational change required to initiate rapid release of the TMD. Overall, our work details the stability of Eag-TMD complexes and the energetic pathway for the dissociation of effector TMDs from their Eag chaperones.

## INTRODUCTION

The ability to secrete proteins into the environment is a central part of bacterial communication and virulence^1^. Secreted proteins often mediate bacterial antagonistic behaviors that shape microbial communities to form complex cooperative and/or competitive networks^2,3^. As such, bacteria have evolved diverse general and specialized secretion mechanisms. These include large membrane spanning complexes designed specifically for the translocation of proteins into the extracellular environment or even directly into rival cells^4–6^. In Gram-negative bacteria, one such complex is the type VI secretion system (T6SS). Generally, the T6SS provides bacteria a competitive advantage in an ecological niche^7^.

The T6SS is a contractile nanomachine that forcibly injects toxic protein effectors into near-by cells in a contact-dependent manner^2,8,9^. The T6SS is structurally related to the T4 bacteriophage tail^10,11^ and consists of a needle, a baseplate, and a membrane spanning complex. The needle is made of hexameric <u>H</u>emolysin <u>c</u>o-regulated <u>p</u>roteins (Hcp) surrounded by a contractile protein sheath that is polymerized starting from the baseplate^10–12^. The membrane-spanning complex orients the needle for ejection into the environment^13,14^. Within the baseplate, a trimeric <u>v</u>aline <u>g</u>lycine <u>r</u>epeat protein G (VgrG) binds the top of the Hcp needle^12,15^. VgrG forms the tip of the T6SS and serves as a modular platform for effector loading^16,17^. VgrG can also be ‘sharpened’ by a proline-alanine-alanine-arginine (PAAR) domain-containing protein^16,18^ or a PIPY domain protein which both bind VgrG to create a structural spike^19–21^. Both PAAR and PIPY domains can serve to load VgrG with cargo effectors^22–26^ and PAAR domains can be covalently linked to an effector to create a specialized or evolved effector^9,27,28^. The T6SS then secretes the entire Hcp-VgrG-PAAR-effector complex. Finally, the effectors inhibit a wide range of essential processes that kill or inhibit the growth of neighbouring prokaryotic and/or eukaryotic cells^6,29^.

A critical step in cargo effector secretion is the selective recruitment or piloting of effectors to the T6SS by chaperone proteins. Chaperones stabilize effectors in a “secretion-competent” state and are essential for the proper delivery and toxicity of numerous T6SS effectors^17,21,27,30^. Several T6SS chaperone and adaptor families have been described, including DUF2169, <u>T</u>ype VI <u>a</u>daptor <u>p</u>roteins (Tap) (DUF4123), and <u>E</u>ffector <u>a</u>ssociated <u>g</u>enes (Eag) (DUF1795). DUF2169 forms complexes with an effector and a PIPY protein to facilitate loading onto VgrG^19,22,31^, Tap adaptors pilot effectors to VgrGs that contain a helix-turn-helix at their C-terminus^17^, and Eag chaperones facilitate the binding of prePAAR containing membrane protein effectors to VgrG^27,32,33^.

The basic architecture of a prePAAR containing effector is an N-terminal prePAAR motif followed by a transmembrane domain (TMD), a PAAR domain, and a C-terminal toxin domain^27^. prePAAR-containing effectors can be further divided into two classes. Class 1 effectors belong to the Rhs family of proteins and possess a single TMD while Class 2 effectors possess two TMD regions directly before and after the PAAR domain^27^. The toxin domains of prePAAR effectors are known to localize to the cytoplasm and inhibit competitor cell growth by acting in the cytoplasm of prey cells, such as by ADP-ribosylation of RNA or protein^34–36^ or rapid depletion of the electron carriers NAD^+^ and NADP^+32,37^. The transmembrane domains (TMDs) are thought to facilitate the translocation of the effector across the prey cell cytoplasmic membrane^30^. Prior to secretion, the TMDs are shielded from the aqueous milieu of the producing cell cytoplasm by their associated Eag chaperone. To promote secretion, Eag chaperones bind the effector TMDs and drastically distort their conformation to inhibit TMD folding^27^. This creates a pre-insertion state to both pilot the prePAAR effector to the T6SS and prevent erroneous membrane insertion before secretion. Additionally, the Eag allows the prePAAR motif to complete a zinc binding motif necessary for the effector PAAR domain to recognize VgrG^27,38^. Notably, Eags are specific for their cognate TMDs and exhibit different effector binding modes.

Given that a bound Eag chaperone would inhibit the proposed membrane penetrating function of the effector TMD, the Eag is presumably removed prior to secretion by the T6SS. Consistent with this hypothesis, Eag chaperones are not secreted alongside their associated effectors^22,23,30,39^. However, there is no current data showing how the Eags are separated from their cognate TMDs. To address this mechanistic question, we took a biophysical and biochemical approach to determine how a T6SS effector TMD is released by an Eag chaperone.

Here, we measure the thermodynamic properties and unfolding kinetics of wild-type Eag-TMD complexes compared to several site-specific point variant Eag-TMD complexes. Our data shows that although Eag-TMDs are remarkably stable, only a small perturbation of the Eag is required to promote TMD dissociation. Specifically, TMD binding to an Eag depends upon a crucial knob-in-hole packing interaction (large against small side chain packing) and/or a hydrogen bond that if removed results in drastically reduced complex stability and rapid effector release. Furthermore, we show that the disruption of these interactions between an Eag and its cognate TMD reduces effector secretion and bacterial fitness, directly correlating our mechanistic findings to biological function. Finally, an Eag-TMD variant X-ray co-crystal complex captures a TMD release intermediate providing key mechanistic details for TMD binding and dissociation from a chaperone. Our work emphasizes the essential role of Eags in TMD folding inhibition and collectively enables us to propose the mechanism for transmembrane domain release from Eag chaperones.

## RESULTS

### Effector TMD binding drastically increases Eag chaperone stability

Given the highly insoluble nature of the TMDs, to date we have been unable to express and purify the TMDs without co-expressing the Eag chaperone. The inability to separately purify Eag and TMD proteins precludes the ability to determine their interaction mechanistic details using common binding assays^40,41^. Instead, we can study the mechanism of effector TMD binding by assaying the stability and thermodynamic properties of Eag-TMD complexes compared to unbound Eags. For this study we use the Eag-TMD complexes SciW-Rhs(1-59) from *Salmonella* Typhimurium and EagT6-Tse6(1- 61) from *Pseudomonas aeruginosa*^27^. For simplicity, the complexes will be referred to as SciW-TMD, EagT6-TMD, or Eag-TMD generally. Notably, the Eag chaperones contain tryptophan residues whereas the TMDs lack tryptophan. As such, by monitoring the change in tryptophan fluorescence during denaturation we can determine how TMD binding affects the energetics and folding behavior of the Eag chaperones.

Using this approach, we first determined the thermostability of the Eag-TMD complexes compared to unbound or apo-Eags. This was done to gauge the stability of the Eag upon TMD binding. When an Eag binds its cognate TMD there is a large increase in the thermostability of the chaperone. Nano differential scanning fluorimetry (nanoDSF) reveals an increase in melting temperature (T_m_) of 30°C for SciW and 14°C for EagT6 (Figure 1A-B). This demonstrates that upon TMD binding the chaperones become as stable as proteins from extremophiles (>65°C)^42^. The purified protein materials used in these experiments are shown in Figure S1.

**Figure 1:**
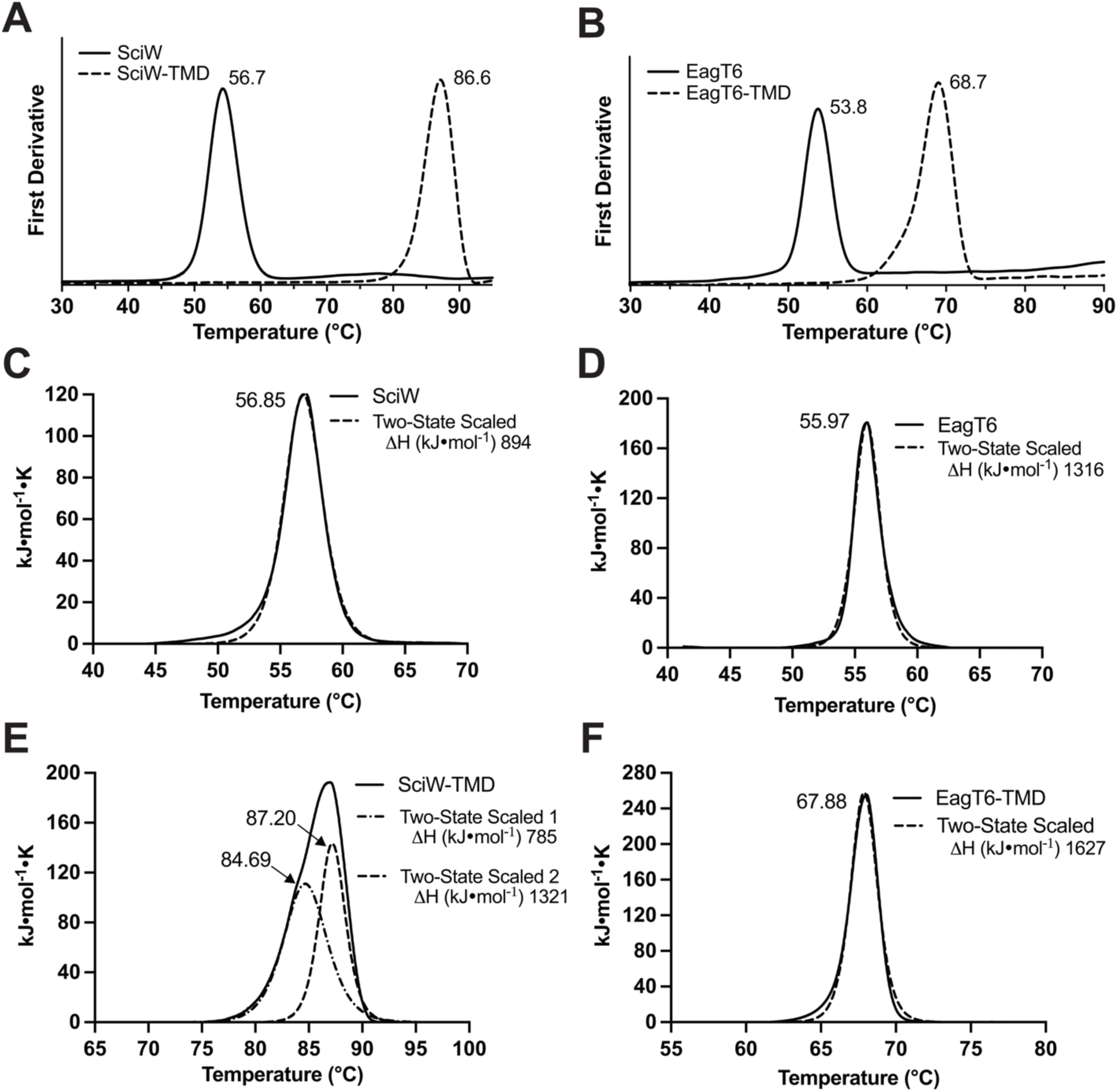
Thermal denaturation of Eag chaperones in complex with effector transmembrane domains. Thermal Denaturation of Eag and Eag-TMD complexes measured by nano Differential Scanning Fluorimetry (nanoDSF). The melting points (T_m_) of **(A)** SciW and **(B)** EagT6 increase to 87°C and 67°C, respectively when bound to their cognate TMDs. Differential scanning calorimetry (DSC) of **(C)** SciW, **(D)** EagT6 **(E)** SciW- TMD and **(F)** EagT6-TMD. Data is fit by a two-state model. All transitions are labeled by fitted T_m_ and enthalpy of unfolding ΔH values are listed for DSC experiments.

As the TMDs are invisible in nanoDSF due to their lack of tryptophan residues, we also compared our results to thermal melting by differential scanning calorimetry (DSC). This was done to observe if the TMDs have melting transitions distinct from their Eag chaperones. The DSC experiments gave nearly identical results to nanoDSF (Figure 1C- F). Furthermore, there were no additional melting transitions in the DSC relative to the nanoDSF for the EagT6-TMD complex indicating that the TMD melts cooperatively with the EagT6 chaperone (Figure 1D). In contrast, the SciW-TMD complex melts in two transitions (Figure 1E). Two transitions observed by DSC but only one by nanoDSF suggests that the TMD changes conformation while still bound to SciW before the complex melts (first transition, Figure 1E). However, in agreement with the nanoDSF data the observed changes in the enthalpy of unfolding (ΔH) for both pairs of Eag chaperones indicates that substantial energy is required to dissociate an Eag-TMD complex.

### Effector TMD binding significantly increases the free energy of Eag chaperone unfolding

We next probed Eag chaperone resistance to chemical denaturation. By measuring Eag and Eag-TMD unfolding by denaturants, we will be able determine the free-energy (ΔG) of Eag stabilization and the binding energy of the Eag-TMD interaction. We denatured Eag and Eag-TMDs with both urea and guanidine hydrochloride (GdmCl) then monitored steady-state Eag unfolding by tryptophan fluorescence. Shown in Figure 2A-B and Table S1, the Eag-TMD complexes are highly resistant to denaturation by urea. Notably, SciW- TMD could not be unfolded with urea (Figure 2A). In contrast, the Eag-TMD complexes could be readily unfolded with GdmCl (Figure 2C-D). The C_50_ (concentration of denaturant at midpoint) values for Figures 2A-D are summarized in Table S1. In agreement with the thermal denaturation data, the chemical denaturation experiments demonstrate significantly increased Eag stability upon TMD binding.

**Figure 2:**
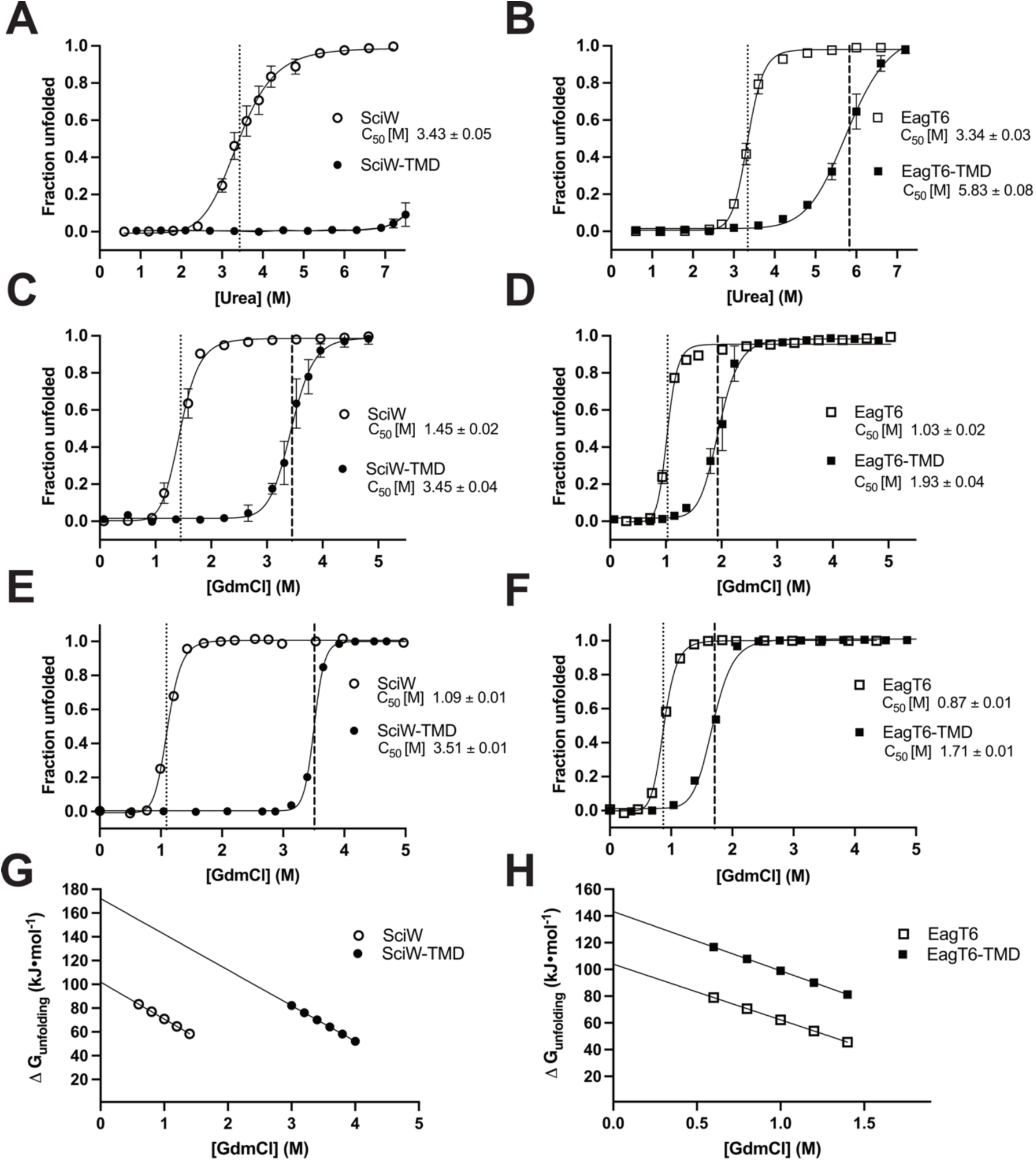
Eag-TMD complexes are resistant to chemical denaturation. Denaturant induced unfolding profiles measured by nanoDSF for EagT6 (○), SciW (□), EagT6-TMD (●) and SciW-TMD (▪) as a function of urea (**A, B**) or guanidine hydrochloride (**C, D**) concentration. (**E, F**) Steady-state unfolding profiles for chaperones as in panels C-D. Data is shown as fraction unfolded. Vertical lines indicate inflection (transition point) at C_50_. (**G, H**) Linear dependence of ΔG_unfolding_ on [GdmCl] of Eag and Eag-TMD complex extrapolated to dilute conditions to obtain ΔG° and the constant of proportionality (m). Results for all parameters are listed in Table 1.

**Table 1:** Unfolding thermodynamic and kinetics of Eag and Eag-TMD complexes.

| Protein | C<sub>50</sub> [M] | $\Delta G^{\circ}_{\text{unfolding}}$<br>(kJ·mol<sup>-1</sup>) | $\ln k^{\circ}_{\text{unfolding}}$ | m<sub>(N-D)</sub> | m' |
| --- | --- | --- | --- | --- | --- |
| **SciW** | 1.09 ± 0.01 | 102.0 ± 1.3 | -5.8 ± 0.4 | -31.1 ± 1.1 | 1.62 ± 0.1 |
| **SciW-TMD** | 3.51 ± 0.01 | 172.2 ± 5.9 | -10.9 ± 0.9 | -30.0 ± 1.7 | 1.57 ± 0.2 |
| **SciW Q58A** | 1.20 ± 0.01 | 104.5 ± 1.4 | -5.0 ± 0.5 | -31.1 ± 1.1 | 1.62 ± 0.1 |
| **SciW-TMD Q58A** | 3.03 ± 0.02 | 157.5 ± 5.0 | -6.7 ± 1.0 | -30.0 ± 1.7 | 1.57 ± 0.2 |
| **SciW L66A** | 1.19 ± 0.01 | 103.5 ± 1.4 | -5.2 ± 0.5 | -31.1 ± 1.1 | 1.62 ± 0.1 |
| **SciW-TMD L66A** | 3.19 ± 0.01 | 163.0 ± 5.3 | -8.9 ± 1.0 | -30.0 ± 1.7 | 1.57 ± 0.2 |
| **EagT6** | 0.87 ± 0.01 | 104.0 ± 2.2 | -1.3 ± 0.2 | -41.7 ± 2.3 | 1.15 ± 0.1 |
| **EagT6-TMD** | 1.71 ± 0.01 | 143.3 ± 2.0 | -2.9 ± 0.4 | -44.3 ± 1.7 | 1.14 ± 0.1 |
| **EagT6 Q58A** | 0.84 ± 0.01 | 103.6 ± 2.1 | 0.08 ± 0.2 | -41.7 ± 2.3 | 1.15 ± 0.1 |
| **EagT6-TMD Q58A** | 1.10 ± 0.01 | 115.4 ± 1.3 | -1.6 ± 0.4 | -44.3 ± 1.7 | 1.14 ± 0.1 |
| **EagT6 L66A** | 0.58 ± 0.01 | 92.7 ± 1.6 | -0.045 ± 0.2 | -41.7 ± 2.3 | 1.15 ± 0.1 |
| **EagT6-TMD L66A** | 0.63 ± 0.01 | 94.9 ± 0.8 | -0.11 ± 0.4 | -41.7 ± 1.7 | 1.14 ± 0.1 |

Now that we have an appropriate system (GdmCl vs. urea), we can determine the free energy of Eag-TMD unfolding. First, we measured the intrinsic fluorescence spectra of the Eags and the Eag-TMDs as a function of increasing GdmCl concentrations (Figure S2). This was done to experimentally find the wavelength maxima (λmax) for the unfolding transitions. Tryptophan fluorescence emission changes from ∼330 nm to ∼350 nm upon protein denaturation, but the exact λmax values can vary per protein which may influence ΔG calculations. Our initial experiments assumed fixed λmax wavelengths of 330/350 nm using a Prometheus to allow for rapid screening^43,44^. Using the data from Figure S2, the fluorescence emission maximum wavelength *λ_max_* as a function of denaturant concentration was determined (Figure S3). If we assume that the Eag chaperones unfold through a two-state mechanism (equation 1), the fraction of tryptophan residues in the unfolded state “α” can be related to the measured *λ_max_*^45^ to plot the transition (equations 2-3). All equation derivations are detailed in methods.

Using equation 2 we have plotted α (fraction unfolded) as a function of denaturant concentration for SciW and EagT6 in their apo and TMD-bound forms (Figure 2E-F and Figure S3). Notably, the experimentally determined *λ_max_* steady-state fluorescence data is in close agreement with the fixed 330/350 nm wavelength data (Figure 2C-D). However, given the slight variation all quantitative thermodynamic and kinetic data will be derived from the experimental *λ_max_* values.

Next, the data in Figure 2E-F was used to calculate the change in unfolding free energy (ΔG_unfolding_) at various denaturant conditions and determine the native ΔG°_unfolding_ at 0 M denaturant (equations 4-6, methods)^46^. From fitting the data in Figure 2E-F to Equation 5, the values of ΔG°_unfolding_ and *m* (slope of the linear extrapolation) for the Eag and Eag-TMD complexes were obtained (Table 1). Figure 2G-H shows the plot of ΔG_unfolding_ as a function of denaturant and the fitted linear extrapolation to 0 M GdmHCl^46^. Both Eags have a similar ΔG°_unfolding_ of ∼103 kJ/mole and each gains significant stability as an Eag-TMD complex. When bound to their cognate TMD, the ΔG°_unfolding_ for SciW increases by ∼71 kJ/mole and the ΔG°_unfolding_ for EagT6 increases by ∼39 kJ/mole. For context, the ΔG°_unfolding_ for lysozyme is ∼30 kJ/mol^47^.

The effector TMD binding energy (ΔG^°^_binding_) and association constant (K_TMD_) can then be obtained from the Eag-TMD ΔG°_unfolding_. If the TMD is assumed to selectively bind the folded form of the Eag with an association constant K_TMD_, the effect of TMD binding upon the standard unfolding free energy for the chaperone can be estimated using equation 7 (methods)^48,49^. ΔG^°^_binding_ is the standard free energy change of TMD binding the chaperone. This directly relates the change in Eag unfolding standard free energy to that of TMD binding at native conditions. The unfolding data of Table 1 allow us to estimate a ΔG^°^_binding_ of approximately -176 kJ/mol for the SciW-TMD complex and a ΔG^°^_binding_ of -116 kJ/mol for the EagT6-TMD complex. Antibodies with pico to femto-Molar (fM, 10^-15^) dissociation constants have a ΔG^°^_binding_ near -100 kJ/mol^50^. The calculated binding constants for the SciW-TMD and EagT6-TMD interactions are on the order of ∼7×10^-32^ M and ∼5×10^-21^ M, respectively. Our thermodynamic data (Figures 1-2) clearly shows that the Eag chaperones bind their TMDs extremely tightly and gain significant stability when in complex with a TMD. This in turn raises the question of the biochemical mechanism needed to overcome the extreme binding energy for TMD release.

### Eag chaperone ɑ-helical packing residue mimics act to inhibit TMD folding

Using the thermodynamic properties of the Eag-TMD complexes combined with their known molecular interactions, we can probe the mechanism of TMD dissociation. Co-crystal structures of SciW-Rhs1 and EagT6-Tse6 show that Eags bind their cognate TMDs by mimicking alpha helical membrane packing^27^. This is accomplished by the Eag dimer folding like a claw around the TMD to create a hydrophobic environment. Additionally, the Eags harbor conserved residues within the pocket of the claw that make hydrophilic bifurcated hydrogen bonds (H-bonds) and knob-in-hole packing interactions with the TMD (Figure 3 and Figure S4A-C). The net effect is to inhibit the TMD from making H-bonds and knob-in-hole packing interactions with itself to properly fold and thus remain bound to the Eag.

**Figure 3:**
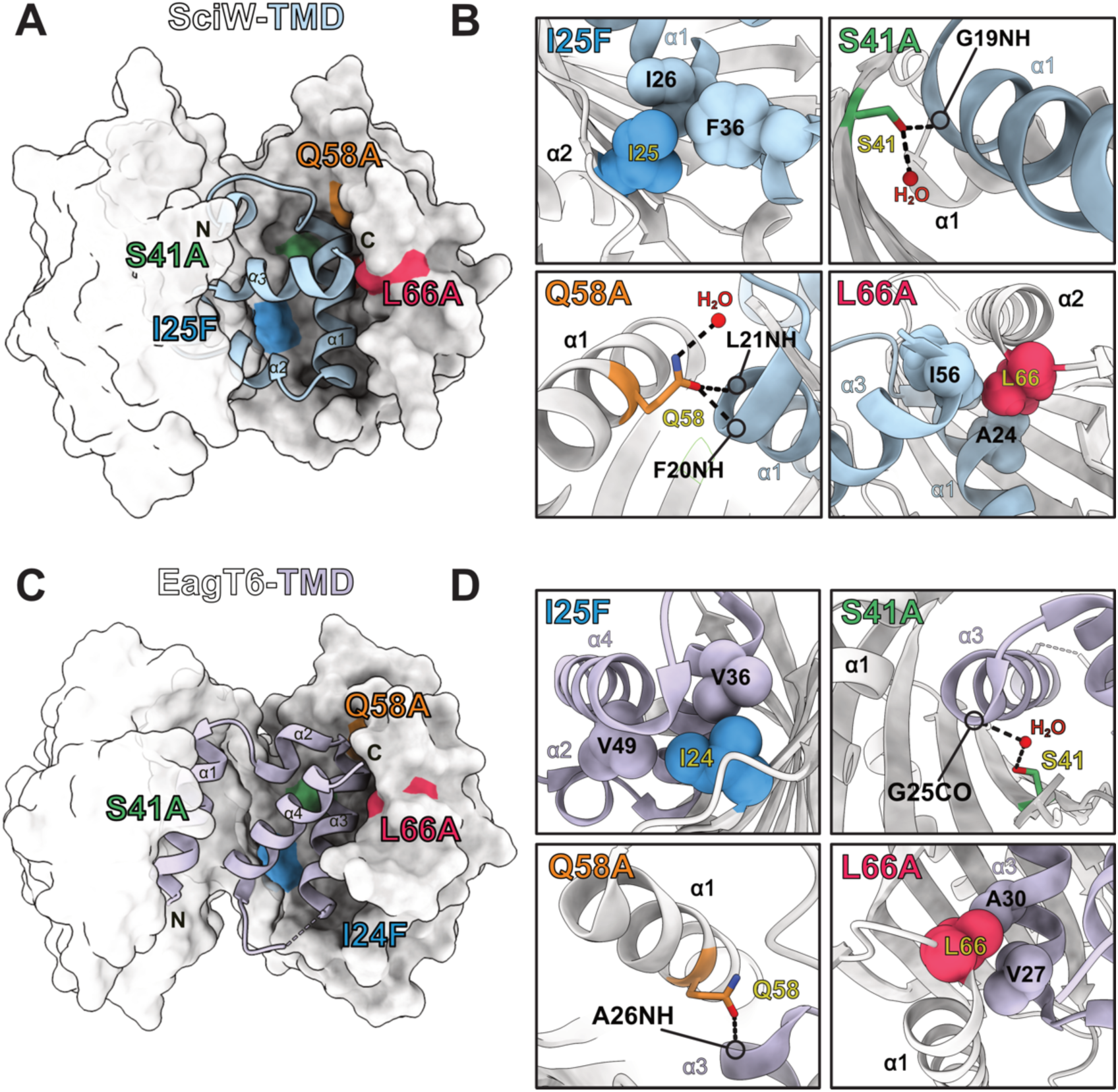
Structural analysis of conserved helical packing interactions between Eag chaperones and TMD helices. Structures of SciW-TMD and Eag-TMD complexes shown with selected point variants by color. **(A)** SciW-Rhs1(1-59) complex showing the position of each point variant. **(B)** Detailed interaction with each SciW point variant and the TMD of Rhs1. **(C)** EagT6-Tse6 (1-61) complex showing the position of each point variant. **(D)** Detailed interaction with each EagT6 point variant and the TMD1 of Tse6. I24/25 is shown in blue and makes hydrophobic packing interactions. S41 is shown in green and makes bifurcated H-bonds (dashed-lines) with the TMD. Q58 is shown in orange and also makes bifurcated H-bonds with the TMD. L66 is shown in red and makes knob-in-hole packing interactions with the TMD. Eag chaperones are shown in light grey and the TMD of Rhs1 in light blue and the TMD1 of Tse6 in light purple. Interactions between the TMD and the second Eag monomer are shown in Figure S4.

To determine which Eag molecular interactions are important for TMD binding and complex stability, we performed site-directed mutagenesis on several conserved residues found in both SciW and EagT6 (Figure 3). The positions of each residue and their specific interactions with the TMD are shown in Figure 3A-B for SciW and Figure 3C-D for EagT6. Note that due to the domain swapped architecture of the dimeric Eag chaperones, these contacts are not equivalent in each Eag monomer. The interaction surfaces of each point-variant in the opposing chain are also depicted in Figure S4D-E.

Eag residue I25/24 was changed to I25F/I24F to interfere with the TMD properly packing against the base of the hydrophobic Eag claw (Figure 3B and D). The Eag hydrophilic residues S41 and Q58 were substituted to alanine because both residues make bifurcated hydrogen bonds to the TMD peptide backbone (Figure 3B and D). Side-chain to main-chain bifurcated H-bonds are crucial for transmembrane helices to properly adopt their tertiary structure^51–53^. EagT6 residue Q102 was also substituted to alanine but attempts to make the same variant in SciW were unsuccessful (Q106) (Figure S4E). The conserved Eag residue L66 was substituted to L66A because it is a mimic of ‘knob-in-hole’ packing interactions. Knob-in-hole packing is when a large hydrophobic residue on one TM-helix acts as a knob to fill the hole created by a small residue such as glycine or alanine on another TM-helix with a GxxxG/A motif^54^. As shown in Figure 3B and D Eag L66 acts as a knob to fill a hole with TMD alanine residues. A shorter side-chain (leucine to alanine) removes the knob. Eag L66 was also of interest because it is on the outer part of the Eag claw. Specifically, the loop containing L66 undergoes a conformational change that moves the Eag claw structure over the TMD upon binding^27^.

After residue substitution, each apo-Eag variant and Eag-TMD variant was purified and its fold assayed by dynamic light scattering (DLS). As shown in Figure S5, all point variants purified similar to wild-type and exhibited the same DLS profile demonstrating that each Eag variant is still dimeric and not aggregated due to misfolding. Furthermore, all Eag-TMD variants still co-purify bound to their cognate TMD showing they are all still functional. Given that we know the ΔG_unfolding_ and ΔG_binding_ for wild-type Eag-TMD (Figure 2), we can now determine how much binding energy is contributed by each Eag residue contact with the TMD and determine mechanistic details of TMD release.

### Eag ɑ-helical packing residue mimics contribute significantly to TMD binding

To probe the effect of Eag variants on TMD binding, we first screened their resistance to chemical denaturation (Figure S6 and Table S2). For SciW-TMD, complex stability was reduced by the residue substitutions but was still significantly more stable than the apo- SciW variants (Figure S6A-D, Table S2). When comparing the change in C_50_ between the apo-EagT6 and TMD bound variants, we observe that all substitutions greatly weaken complex stability (Figure S6E-I, Table S2). However, EagT6 I24F, L66A, and Q102A appear to have the greatest effect (small or no change in bound to unbound variant C_50_). This shows that EagT6-TMD binding is severely weakened by either the loss of knob-in-hole (L66A) or bifurcated H-bond interactions (Q58A and Q102A) (Figure 3, Figure S4). Additionally, steric hindrance of proper TMD packing by a larger hydrophobic residue (I24F) is also severely disruptive. For the SciW-TMD complex the loss of bifurcated H-bonds (Q58) or knob-in-holes (L66A) is substantial but less pronounced (Figure S6 and Table S2).

The variants Q58A and L66A represent two distinct modes of ɑ-helix membrane protein packing and affect both SciW-TMD and EagT6-TMD complex stability. Given this, we selected these variants for ΔG_unfolding_ quantification by intrinsic protein fluorescence spectral analysis. The tryptophan fluorescence wavelength maxima (*λ_max_*) for the Q58A and L66A point variants were determined and used to plot α (fraction unfolded) as a function of denaturant concentration using equation 6 (Figure S7, Figure 4A-D, and methods). The ΔG_unfolding_ for the variants was calculated from fitting α (fraction unfolded) with equations 1-6 and ΔG^°^_binding_ from equation 7 similar to the wild-type. Because the “m” parameter in equation 6 only depends upon the change in surface accessible area upon unfolding^55^, the α (fraction unfolded) values were fitted globally sharing the “m” parameter in these subsets: 1) apo-SciW variants, 2) SciW-TMD variants, 3) apo-EagT6 variants, 4) EagT6-TMD variants. The resulting parameters from the fits are listed in Table 1. The dependence of ΔG_unfolding_ on denaturant concentration with fitted extrapolations to 0 M denaturant (ΔG°_unfolding_) are shown in Figure 4E-F.

**Figure 4:**
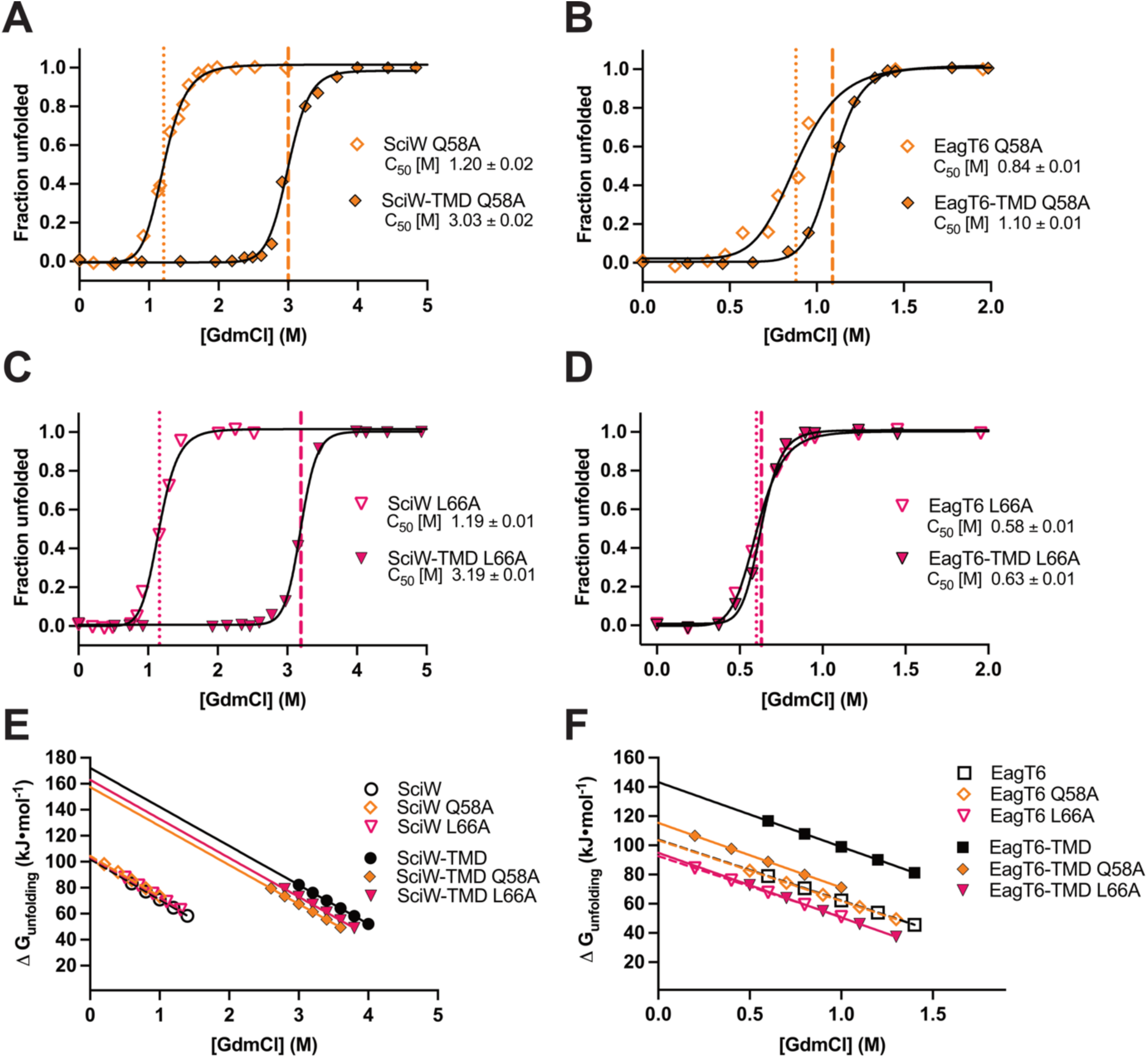
Eag-TMD point variants show reduced complex stability by chemical denaturation. Denaturant induced unfolding profiles of variant Eag and Eag-TMD complexes. (**A, B**) Q58A variant unfolding profiles. (**C, D**) L66A variants unfolding profiles. **(E, F)** Linear dependence of ΔG_unfolding_ on [GdmCl] of Eag and Eag-TMD variants extrapolated to dilute conditions to obtain ΔG°_unfolding_. Eag Q58A variants are plotted as an orange open diamond. Eag-TMD Q58A variants are plotted as an orange filled diamond. Eag L66A variants are plotted as a fuchsia open triangle. Eag-TMD L66A variants are plotted as a fuchsia filled triangle. Wild-type SciW and SciW-TMD complexes are shown as open and black circles respectively. Wild-type EagT6 and EagT6-TMD complexes are shown as open and black squares respectively. Wild-type fits are replotted from Figure 2 to compare to variants. The results of all fits are shown in Table 1.

Compared with the wild-type ΔG°_unfolding_, the point variants Q58A and L66A have little effect on the ΔG°_unfolding_ for apo-SciW. However, both residue substitutions lowered the ΔG°_unfolding_ of the SciW-TMD complex indicating a reduction in TMD binding affinity (Table 1 and Figure 4A,C,E). The ΔG^°^_binding_ values for SciW Q58A and L66A with the TMD are -142 kJ/mol and -154 kJ/mol, respectively, compared to -176 kJ/mol for wild-type. We can conclude that the bifurcated H-bonds provided by Q58 contribute ∼34 kJ/mol to TMD binding, and the knob-in-hole interactions of L66 contribute ∼22 kJ/mol. Within experimental error, these are roughly 15-20% of the total SciW-TMD binding energy. This also increases the dissociation constant for Q58A and L66A relative to wild-type by several orders of magnitude to ∼1×10^-25^ M and ∼7×10^-28^ M, respectively.

For EagT6, the point variants Q58A and L66A have a more pronounced effect on complex stability and TMD binding. The folding free energy of apo-EagT6 is minimally affected by the Q58A substitution, but the stability of EagT6 is reduced by L66A (Table 1 and Figure 4B,D,E). However, the EagT6 L66A variant chaperone is still properly folded and functional as it co-purifies bound to the TMD (Figure S5). Moreover, we observe drastic reductions in the ΔG°_unfolding_ for the EagT6-TMD variant complexes showing that Q58 and L66 play critical roles in TMD binding. For the Q58A, the ΔG^°^_binding_ to the TMD has reduced in magnitude from -116 kJ/mol to -60 kJ/mol. Thus, we can conclude that the bifurcated H-bonds and packing interactions of Q58 with the TMD contribute at least 56 kJ/mol or about 1/2 the binding energy. This increases the dissociation constant to ∼3x10^-11^ M or about 10 orders of magnitude. For EagT6 L66A, the value of ΔG°_unfolding_ is similar for the variant apo and TMD bound. This indicates that minimal TMD binding to the EagT6 L66A variant exists under our assay conditions. Our fluorescence experiments are performed under dilute conditions (∼ 1 μM chaperone) and in the absence of crowding agents, it is likely that the TMD has dissociated prior to the assay. This shows that the binding constant K_TMD_ for the L66A substitution has been reduced to less than 10^6^ M^-1^ putting an upper limit of ∼ -35 kJ/mol upon the ΔG^°^_binding_. Our data shows that the knob-in-hole packing of residue L66 is crucial for maintaining the EagT6-TMD complex.

### Minor variations to the Eag molecular surface accelerate the rate of TMD release

Our biophysical data thus far has been measured at equilibrium conditions. However, T6SS assembly and firing is a dynamic process that can occur within several seconds^56–58^. Given this, how quickly the TMD can be released prior to secretion is of central importance to the function of the Eag chaperone. To determine the rate of TMD release, we investigated the unfolding kinetics of wild-type and variant Eag chaperones using stopped-flow spectrofluorimetry. Unfolding fluorescence trace data as a function of time for wild-type Eag and Eag-TMD complexes is shown in Figure S8 and point variant traces in Figure S9. The relaxation profiles can be described with a mono-exponential decay and the native unfolding rate constants (k°_unfolding_) calculated using chevron plot analysis by extrapolation to 0 M GdmCl^46^ (Figure 5 and Table 1, equations 8-9, methods).

**Figure 5:**
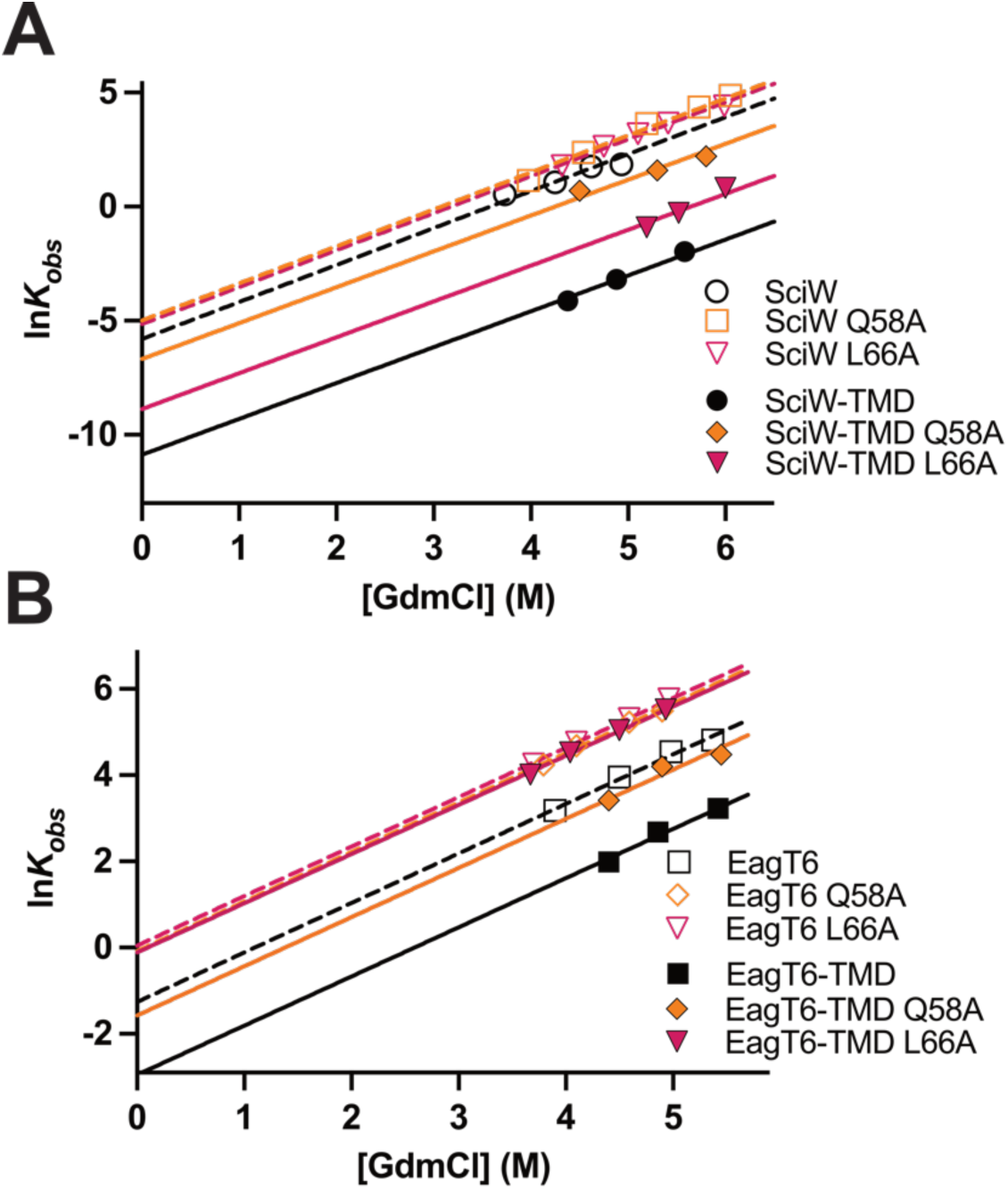
Unfolding kinetics of wild-type and point-variant Eag chaperones. The observed unfolding rates (kobs) of **(A)** SciW and SciW-TMD and **(B)** EagT6 and EagT6- TMD as function of GdmCl concentration. Data was linearly extrapolated to 0 M GdmCl to obtain the rate of unfolding in buffer (ku). Fit parameters (mu) and lnku for the linearly regression are shown in Table 1. Stopped-flow traces are shown in Figures S8-9.

Wild-type SciW and the Q58A and L66A variants have unfolding rate constants (k°_unfolding_) that are indistinguishable within the limits of experimental uncertainty (Figure 5A and Table 1). The k°_unfolding_ constants reveal that the unfolding/folding lifetime for SciW is on the order of ∼5 minutes. Again, this highlights that the variants have no effect on the biophysical properties of the SciW chaperone itself. However, both Q58A and L66A significantly increase the rate of unfolding/folding for the SciW-TMD complex (Figure 5A). The wild-type SciW-TMD complex has a lifetime on the order of ∼15 hours but is reduced to ∼2hrs for SciW L66A and ∼16 minutes for SciW Q58A. The disruption of a knob-in-hole (L66) or bifurcated H-bond (Q58) increases the rate of TMD release by 7.5x and 60x respectively, but only at the cost of 15-20% binding energy (Figure 4E, Table 1).

The wild-type EagT6 chaperone has an unfolded/folded lifetime of ∼5 seconds which increases to ∼18 seconds when bound to TMD1 of Tse6 (Figure 5B and Table 1). The extreme difference in the folded lifetime of apo EagT6 compared to apo SciW was unexpected, as both chaperones have the same ΔG°_unfolding_ (Table 1) and domain swapped dimeric structure (Figure 3)^27,59^. Additionally, we find that both Q58A and L66A increase the rate of unfolding/folding for the apo EagT6 chaperone to ∼1 second. Interestingly, this differs from the steady-state data where EagT6 Q58A is indistinguishable from wild-type. However, similar to SciW both EagT6 Q58A and EagT6 L66A greatly increase the rate of EagT6-TMD complex unfolding/folding. EagT6 Q58A reduces the lifetime of the EagT6-TMD complex to ∼5 seconds. This is ∼4 second difference in lifetime for the Q58A variant as compared to ∼13 seconds for wild-type, demonstrating that the increased TMD release rate is not primarily due to the faster unfolding of the apo-variant. For EagT6 L66A we observe no kinetic difference with the TMD (Figure 5B and Table 1). Again, this is likely because TMD release by EagT6 L66A is faster than we can measure. In agreement with the steady-state data, the knob-in-hole interactions provided by EagT6 L66 are essential to keep the TMD bound to the Eag.

### The transition state intermediate of TMD release from Eags differs by effector class

The effect of Eag variants upon the unfolding rate (k°_unfolding_) and energy (ΔG°_unfolding_) also reveals mechanistic details about the TMD dissociation transition state. By applying a “φ- value analysis” to the measured Eag thermodynamic and kinetic parameters we gain insight into the rate-limiting step of the unfolding transition state. The φ-value is the ratio in change of free energy for unfolding/folding kinetics (ΔG^‡^_unfolding_ = -RT(lnk°_unfolding_)) to the equilibrium free energy of unfolding/folding (ΔG°_unfolding_) between wild-type and variants (equation 10, Table 1). This gives a φ-value between 0 and 1, where 0 represents a dynamic or unstructured transition state and 1 an ordered or structured transition state^60^. The φ-value analysis yields 0.7 ± 0.4 for SciW Q58A, 0.5 ± 0.2 for SciW L66A, and 0.1 ± 0.3 for EagT6 Q58A. A φ-value calculation for EagT6 L66A was not possible because the TMD had already dissociated. The small φ-value for EagT6 Q58A indicates a dynamic or disordered conformation at the unfolding transition state of the EagT6-TMD complex. This could be due to that the TMD is conformationally dynamic at the transition state and/or that the TMD has already been released. Moreover, release of the TMD from EagT6 and the unfolding of EagT6 are uncoupled processes. Given the inability to determine a φ-value for EagT6 L66A a similar disordered transition state is likely. For SciW Q58A and L66A, the φ-values are both 0.5 or higher, which is more characteristic of a structured unfolding transition state. Additionally, the φ-values suggest that the TMD is bound to SciW during the unfolding transition state. This indicates that the release of the TMD and unfolding of SciW are coupled. Overall, our biophysical characterization shows that the exact mechanistic details and transition state of TMD release differ between SciW (class 1 effector TMDs) and EagT6 (class 2 effector TMDs). First, that disruption of the H-bond or knob-in-hole provided by Q58 or L66 is sufficient to release the TMD from EagT6, but TMD release from SciW requires disruption of more than one contact. Second, that the release of the TMD from SciW involves a stable conformation intermediate of either the TMD, SciW or both.

### EagT6-TMD stability variants inhibit effector secretion and decrease bacterial fitness

To observe how our biophysical characterization relates to biological activity, we next determined the contribution of our Eag point variants to the function of the chaperones *in vivo*. We performed growth competition assays using the chaperone-effector pair EagT6- Tse6 in *P. aeruginosa*. EagT6-Tse6 was chosen for our model as our point variants have a significantly greater effect on Tse6 TMD release from EagT6 as compared to SciW and the TMD from Rhs1 (Figures 4-5). For example, in EagT6 L66A is sufficient to rapidly release the TMD *in vitro*. Additionally, the T6SS of *P. aeruginosa* is a highly tractable model system since deletion of the negative T6SS regulator *retS* results in strains with constitutive T6SS activity^17,30,32^.

Competition assays were performed with a *P. aeruginosa* donor and an isogenic Tse6-susceptible recipient that lacks Tse6 and the immunity protein Tsei6 (*Δtse6Δtsei6*)^27,30^. The donor strains encode the wild-type EagT6 chaperone (parent), no chaperone (*ΔeagT6)* or one of the EagT6 point variants expressed from their native chromosomal locus. When co-cultured on a solid surface, the wild-type donor readily outcompetes the recipient strain (Figure 6A). Consistent with our biophysical data, any of the *eagT6* alleles encoding variants of the chaperone display attenuated fitness against the Tse6-susceptible recipient. Specifically, EagT6 L66A and Q102A most strongly contribute to the function of EagT6 *in vivo* and therefore the proper secretion of Tse6. This agrees with our biophysical data that L66 is crucial to EagT6-TMD complex stability. Since Tse6 requires EagT6 binding for cytosolic accumulation in *P. aeruginosa*^30^, we reasoned that Tse6 levels may be reduced in strains harboring our mutant *eagT6* alleles. Indeed, expression of each EagT6 variant decreased intracellular Tse6 levels (Figure 6B). Furthermore, the relative abundance of Tse6 in the EagT6 variant backgrounds correlates with the contribution of that residue to TMD binding energy (Figures 4-6, Figure S6, and Table 1). Overall, these findings demonstrate that the knob-in-hole packing provided by EagT6 L66 with the TMDs of Tse6 is critical for the stability of the EagT6-Tse6 complex *in vivo* and proper secretion of the effector.

**Figure 6:**
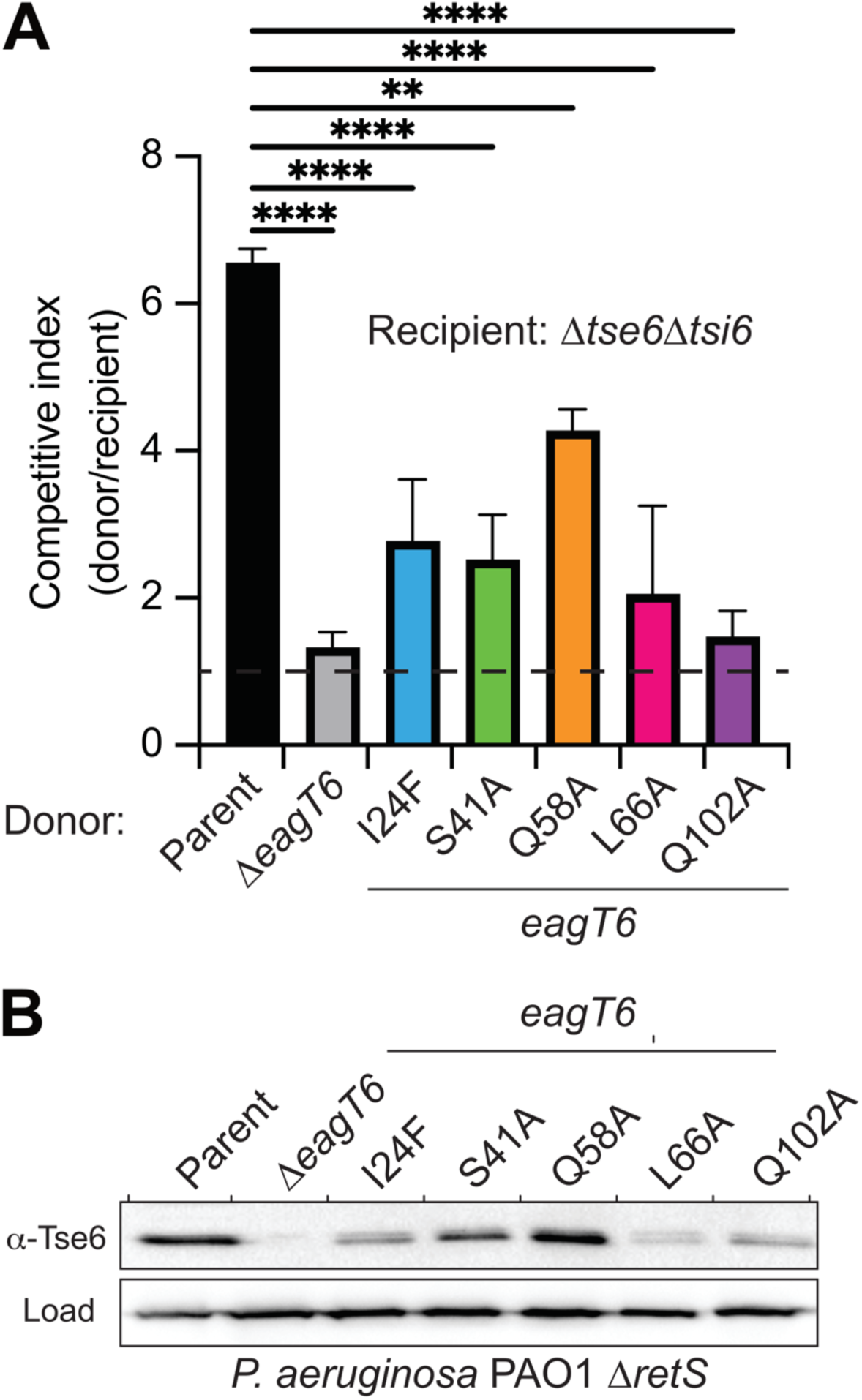
Intraspecific competition experiments between *P. aeruginosa* PAO1 predator and prey strains. **(A)** Bacterial competition experiment between strains harboring wild-type EagT6 and EagT6 point variants. The competitive index is calculated as the change (final/initial) in the donor to recipient ratio. The parent is PAO1 *ΔretS.* Horizontal dashed line at a competitive index of 1 indicates the line of no competitive advantage. Asterisks indicate statistically significant differences, error bars represent SDM, n=3. **(B)** Western blot analysis of Tse6 expression in the indicated *P. aeruginosa* PAO1 strains. Parent is PAO1 *ΔretS.* RNA polymerase was used as a loading control.

### A crystal structure of SciW-TMD L66A suggests a TMD release intermediate

To further probe the molecular details of Eag-TMD complex stability we attempted to crystallize a point-variant Eag-TMD complex. We were able to obtain diffracting crystals of SciW-TMD L66A and obtain phases by molecular replacement (Figure 7). Two SciW- TMD complexes were present in the asymmetric unit and all chains exhibited very high B-factors suggesting conformational dynamics (Figure S10A and Table 2). Specifically, one SciW-TMD L66A complex (chains D-F) showed several areas of missing density and the TMD was not confidently modelled. However, the other SciW-TMD L66A complex (chains A-C) showed strong density for the TMD (Figure S10B). Strikingly, the conformation of the bound TMD is significantly different from the wild-type complex (Figure 7 and Figure 3A). Given the high B-factors, lack of strong TMD density for one complex, and the observed alternate TMD conformation, the captured SciW-TMD L66A structure likely represents a TMD release intermediate.

**Figure 7:**
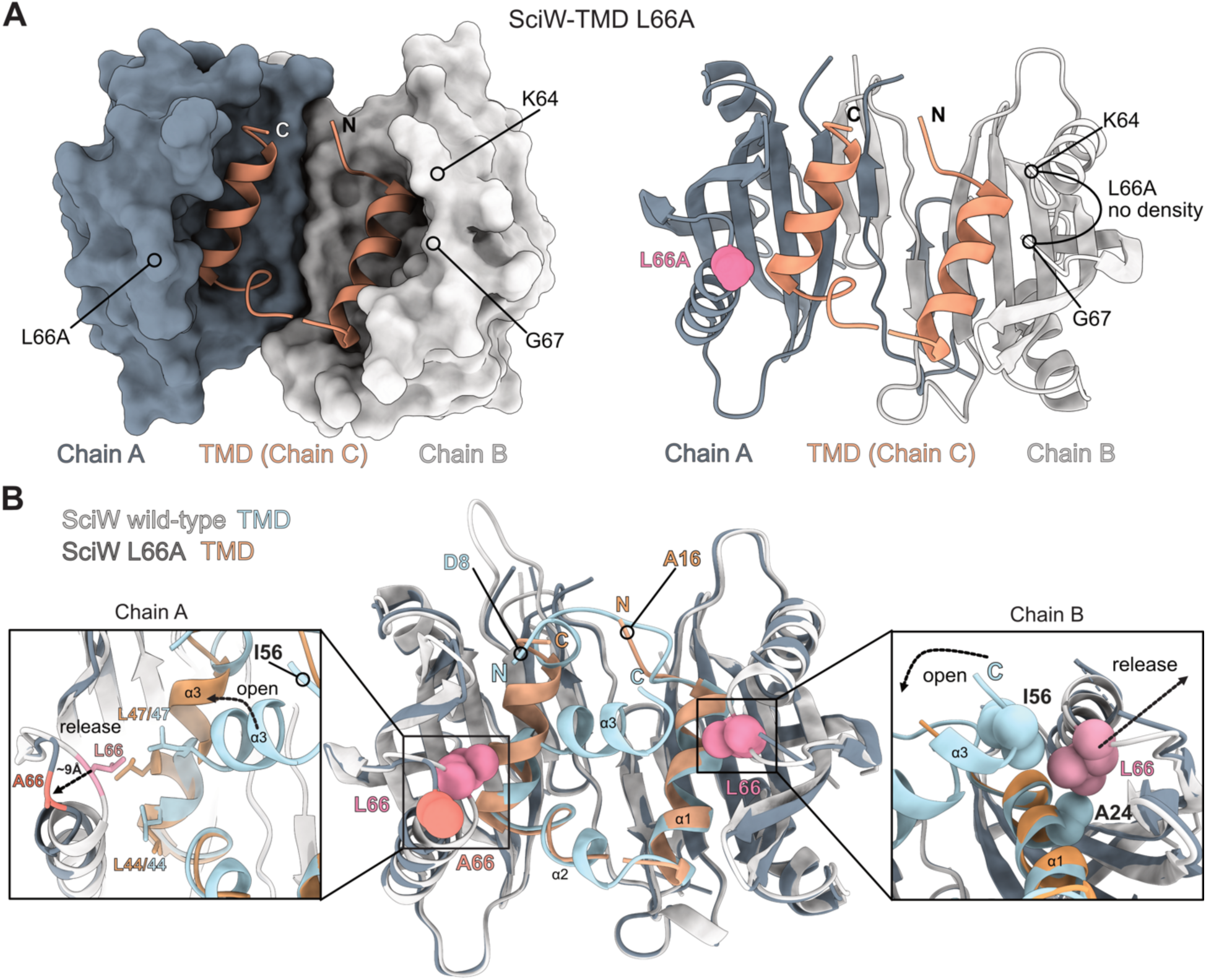
An X-ray crystal structure of SciW-TMD L66A captures an TMD release intermediate. **(A)** Structure of chains A-C. SciW is shown as a molecular surface and the TMD of Rhs1 in dark salmon (left). Cartoon ribbon diagram of chains A-C with the position of L66A indicated by space-filing in pink (right). The unmodelled loop region containing L66A in chain B is also indicated. **(B)** Structural comparison of wild-type SciW-TMD (grey- light blue) and SciW-TMD L66A (dark grey-dark salmon). Boxes show a zoom-in of the conformational changes due to the L66A variation at chain A (left) and chain B (right). Wild-type L66 is shown in dark red. L66A (A66) is shown in orange-red. Residues at the N-terminus of the SciW-TMD and SciW-TMD L66A complexes are indicated.

**Table 2:** Data collection and refinement statistics for SciW-TMD L66A.

|  |  |
| --- | --- |
| **Data Collection** | SciW-Rhs1(1-59) L66A |
| Wavelength (Å) | 1.1808 |
| Space group | C 1 2 1 |
| Cell dimensions |  |
| a, b, c (Å) | 90.67 52.02 177.45 |
| $\alpha, \beta, \gamma$ (°) | 90 99.76 90 |
| Resolution (Å) | 45.16 - 3.05 (3.26 - 3.05) |
| Total reflections | 104432 (19002) |
| Unique reflections | 15847 (2884) |
| CC (1/2) | 0.996 (0.492) |
| $R_{merge}$ | 0.151 (1.948) |
| $R_{pim}$ | 0.064 (0.816) |
| $I/\sigma$ | 8.2 (0.9) |
| Completeness (%) | 99.8 (100.00) |
| Redundancy | 6.6 (6.6) |
| **Refinement** |  |
| $R_{work} / R_{free}$ (%) | 26.1 / 29.8 |
| Average B-factors | 175.8 |
| Protein | 176.0 |
| Water | 143.5 |
| No. atoms |  |
| Protein | 4877 |
| Water | 38 |
| Rms deviations |  |
| Bond lengths (Å) | 0.002 |
| Bonds angles (°) | 0.63 |
| Ramachandran plot (%) |  |
| Total favoured | 95.33 |
| Total allowed | 4.34 |
| PDB code | 9MVG |

Statistics for the highest resolution shell are shown in parenthesis

SciW-TMD L66A retains a dimeric structure similar to wild-type however the loop containing L66A at the tip of each claw is in a different conformation (Figure 7A, Figure 3). There is no observable electron density for the loop between SciW residues K64 and G67 of chain B (Figure 7A right). Furthermore, the TMD is also no longer in a twisted asymmetric conformation. In wild-type SciW, the TMD (blue) adopts an asymmetric binding mode making non-equivalent contacts with the chaperone dimer (Figure 3, Figure S4, and Figure 7B)^27^. Instead, the TMD folds as two symmetric helices packed similarly into the bottom of the L66A chaperone claw (Figure 7A).

When comparing the SciW-TMD L66A structure to the wild-type complex we observe that the L66A variant is unable to form critical knob-in-hole and hydrophobic packing interactions with the TMD (Figure 7B). First, L66A in chain B no longer forms a knob-in-hole with Rhs1-TMD A24 in helix ɑ1 or packs against residue I56 in helix ɑ3 due to the shorter sidechain of the substituted alanine. This in turn frees SciW loop K64 to G67 and releases the C-terminus of the TMD from the chaperone, which causes the TMD to open outwards from its normal twisted conformation (Figure 7B, chain B). Simultaneously in chain A, L66A no longer packs against TMD residues L44 or L47 in helix ɑ3. The loop containing A66 moves ∼9Å away also releasing the TMD to open in a hinge-like fashion forming an ɑ-helix that fills the chaperone binding pocket. This is in part due to TMD residue L47 moving into the space that the SciW L66 side chain previously occupied. Additionally, the bound N-terminal prePAAR region (residues 8-15) of the TMD is pushed out and released from SciW. A movie demonstrating the conformational change is provided (Movie S1). Our structural data shows why L66 is important for SciW-TMD complex stability and in agreement with our φ-value analysis, suggests a partially ordered transition state during TMD dissociation. Given that residue L66 is critical for EagT6-TMD complex stability and that L66A allows the visualization of SciW-TMD unfolding intermediate, a conformational change to move L66 away from the TMD is likely central to the mechanism of TMD release for both classes of Eag chaperones.

## DISCUSSION

Here, we use a combination of biophysical, molecular genetics, and structural biology methods to investigate the mechanism of effector transmembrane (TMD) binding to Eag chaperones. Our work shows that the Eag chaperones SciW and EagT6 exhibit an extremely strong binding affinity (tighter than antibody-antigens) for the TMDs of their cognate effectors Rhs1 and Tse6 (Figures 1-2). As chaperones sequester TMDs by mimicking alpha-helical membrane packing, we designed point variants to disrupt these contacts and probe how the TMD might dissociate from an Eag (Figure 3). Our quantitative thermodynamic and kinetic measurements demonstrate that the disruption of bifurcated hydrogen bonds (Q58) or knob-in-hole packing interactions (L66) between the Eag and TMD is sufficient to cause EagT6 to release the TMD both *in vitro* and *in vivo* (Figures 4-6). Additionally, a co-crystal structure of SciW-Rhs1 L66A reveals an intermediate TMD-release state. The structure shows that the loss of key knob-in-hole packing interactions drastically changes the conformation of the TMD within the Eag and increases the dynamics of the complex (Figure 7). Given that only small perturbances in Eag-TMD complexes drastically lowers the binding energy and leads to a large conformational change in the TMD, our work suggests that the mechanism of TMD release from an Eag only requires a subtle conformational change to disrupt one or two key Eag-TMD contacts.

Given the sum of our data, we propose the following model to catalyze TMD release. First, the Eag chaperone binds the TMD of its cognate effector to pilot the effector to VgrG and catalyzes completion of the PAAR domain with the prePAAR motif^27^. Notably, this removes the prePAAR interactions from the Eag. However, this is not enough to dissociate the Eag as Eag-effector-VgrG complexes can be readily isolated and studied by cryoEM^27,30,36^. Next, we hypothesize that the Eag is removed from the TMD within the baseplate/membrane complex. Again, this is because Eag-effector-VgrG complexes can be readily isolated from Gram-negative bacteria which strongly suggests that the dissociation catalyst is not cytoplasmic but specific to the T6SS apparatus. Within the baseplate, a conformational change in Eag is induced to disrupt the Eag-TMD complex. Our data indicates that the induced conformational change moves L66 away from the TMD. Residue L66 is critical to Eag-TMD complex stability and undergoes a conformational change to initially bind the TMD^27^. Disruption of the critical knob-in-hole and packing interactions provided by L66 lowers the incredibly high Eag-TMD binding energy and increases the rate of TMD dissociation. This in turn makes the Eag release the TMD, albeit SciW-TMD (class 1) and EagT6-TMD (class 2) proceed through different folding transition state intermediates. Unlike EagT6, our data shows that release of the TMD from SciW requires more than just disruption of L66. Residue Q58 is part of the ɑ- helix that ends with the L66 loop, which also moves to accommodate TMD binding. This suggests that a conformational change that moves L66 would also displace Q58 (and other contacts) and therefore may allow TMD dissociation of either effector class. Alternatively, the conformation seen in the L66A variant occurs first creating a short-lived intermediate followed by disruption of other contacts. Currently it remains to be discovered what protein or other factor catalyzes and provides energy for the chaperone conformational change. Regardless, our biophysical data provides molecular details for the energetic mechanism TMD release.

Although Eag-TMD complex stability can be overcome by disruption of a knob-in-hole or H-bond interaction, it is unknown what biological factor catalyzes TMD release. Given the extremely high TMD binding energy there may be a protein that provides energy to dissociate the TMD. One possibility is that a T6SS structural component of the baseplate or membrane complex such as TssK, TssL, or TssM binds the Eag chaperone^61,62^. This binding could then induce the conformational change in the Eag as described by our biophysical data. TssM possess ATPase activity which phosphorylates TssL as part of Hcp recruitment^63^ and could provide energy to dissociate the TMD. However, our biophysical data clearly demonstrates that the energetics of TMD release only requires outcompeting one Eag residue interaction (L66), at least for EagT6. Residue L66 is on a solvent accessible loop on the very end of the Eag helix that changes conformation to bind the TMD. Given this, a T6SS protein only needs to bind the Eag and outcompete L66 to lower the energy of TMD binding and allow dissociation. Finally, the T6SS internal apparatus is shielded from the bulk solvent^61^. This could result in an environment that alters Eag sidechain pKas disrupting bifurcated hydrogen bonds or outcompete the van der Waal knob-in-hole interactions with the TMD. Regardless, additional experimentation is required to determine if a T6SS protein induces Eag chaperone removal from the effector.

Our biophysical data also provides additional insight to understand Eag-TMD dissociation in the timeframe of T6SS assembly. The EagT6-TMD and SciW-TMD complexes have lifetimes on the order of ∼18s and ∼15hrs respectively. Removal of the knob-in-hole and packing interactions of L66 reduces the TMD-bound lifetime to ∼2hrs for SciW and release of the TMD is unmeasurably fast for EagT6 (Figure 5 and Table 1). Given that the T6SS can assemble and fire on the order of seconds to minutes^56–58^ our biophysical data fits well for EagT6-TMD within the known time frame of T6SS kinetics. Additionally, our data supports that the Tse6 TMD release mechanism is two-state. In contrast, the release of the TMD from SciW is too slow to depend only on the removal of one knob-in-hole interaction. The SciW Q58 variant accelerates complex dissociation to ∼16 minutes also supporting that release of the TMD from SciW requires a more substantive disruption than EagT6 as proposed in our biophysical model. Alternatively, this may also suggest a multi-step mechanism for TMD release from SciW. This hypothesis is supported by our SciW-TMD L66A crystal structure that captures a TMD release intermediate and the DSC data that shows a thermal unfolding intermediate for the TMD. However, our chemical unfolding data for SciW is readily modelled by a 2-state mechanism suggesting that further kinetic experiments are required to resolve if other SciW-TMD unfolding intermediates exist. However, our data strongly supports that for both chaperones release of the TMD includes a conformational change to displace L66.

Based on our data, the precise mechanistic details for TMD release differ between SciW-TMD and EagT6-TMD. In SciW the TMD adopts a twisted conformation compared to an ordered anti-parallel helix found in EagT6^27^. These structural differences likely affect the mechanism of TMD dissociation. For EagT6-TMD, a conformational change to disrupt L66 is sufficient to displace the anti-parallel TMD helix. For SciW-TMD, the higher complex stability and TMD intermediate observed by our biophysical data may act as an energetic guardrail. Such an energetic guardrail may have biological relevance directly related to the structure of class 1 and class 2 prePAAR effectors^27^. EagT6 binds a class 2 effector that has 2 TMD regions. This requires 2 Eags to stabilize the TMDs and prevent their membrane-penetrative conformation. In contrast, SciW binds a class 1 effector that has only 1 TMD region which would not require the extra energy barriers to remove a second Eag then complex with another TMD. Given this, it is likely that class 1 complexes have evolved higher stability than class 2 complexes to assure inhibition of TMD folding. Overall, our work has provided a detailed biophysical description of the energetics of Eag-TMD complex stability, Eag TMD effector class specificity, and the mechanism for TMD release.

## MATERIALS AND METHODS

### Molecular cloning of protein expression constructs

Wild-type Eag chaperones SciW from *Salmonella* Typhimurium (Eag: SL1344_0268) and EagT6 from *Pseudomonas aeruginosa* (Eag: PA00923) and their respective point variants were commercially synthesized from Twist Bioscience in the vector pET-29b(+). A C-terminal VSV-G tag for detection by western blot was added using restriction sites NdeI and Xho1. Eag-TMD complexes SciW-Rhs1_NT_ (Eag: SL1344_0286, Rhs1 (1-59)_TMD_ :SL1344_0285) and EagT6-Tse6_NT_ (Eag: PA0093, Tse6 (1-61)_TMD_: PA0094), were created previously in the vector pETDUET-1 in Ahmad et. al.^27^. C-terminal VSV-G-tagged Eag chaperones were both cloned into multiple cloning site 2 (Nde1/Xho1) and N-terminal domains of the effectors were cloned into multiple cloning site 1 with a C-terminal His_6_- tagged (Nco1/HindIII). Eag-TMD point variants were created using Q5 Site-directed mutagenesis kit (NewEngland Biolabs, NEB) by amplifying the full-length construct using primer pairs outlined in Table S5. Primers were generated from Integrated DNA Technologies (IDT) and constructs were confirmed by sanger sequencing (SickKids Toronto).

### Protein expression and purification of Eag chaperones

Eag chaperones and their point variants were over expressed in *E. coli* BL21(DE3)-Gold cells using Lysogeny Broth (LB) culture medium and supplemented with 50 μg/mL kanamycin. Cells were grown at 37 °C in a shaking incubator overnight and the next day diluted in 3 liters of LB broth and grown to OD600 of 0.6. Protein expression was induced by the addition of 1 mM isopropyl β-d-1-thiogalactopyranoside (IPTG) and cells were further incubated for 20 hours overnight at 20 °C. Next, the cells were harvested by centrifugation at 4200*g* for 30 minutes and resuspended in Eag lysis buffer (50 mM Tris pH 8). PMSF and MgCl_2_ were added at final concentrations of 1 mM and 10 mM respectively, along with a small amount of DNase. The cells were then lysed using an Emulsiflex-C3 High Pressure Homogenizer (Avestin) and centrifuged at 17,000*g* for 25 minutes at 4 °C. The supernatant was passed over a Q-sepharose gravity column (GE Healthcare) equilibrated with 2 x column volumes of Eag lysis buffer for purification by ion exchange. The column was washed with 3 x column volumes of lysis buffer before being subjected to a stepwise gradient of 1 x column volumes of Eag lysis buffer containing increasing concentrations of NaCl as follows: 50 mM NaCl, 100 mM NaCl, or 1M NaCl. Fractions from the 50 mM elution step (50 mM Tris pH 8, 50 mM NaCl) were concentrated to 2 mL by centrifugation using a 3 kDa concentrator (Amnicon). The concentration material was further purified by size-exclusion chromatography (SEC) using a HiLoad (16/600) Superdex75 column (Cytvia) with an AKTApure (Cytvia). The SEC column was equilibrated with gel filtration buffer (50 mM Tris pH 8, 250 mM NaCl, 1mM 2- Mercaptoethanol (BME) before samples were injected. All SEC purifications were performed at 4*°*C. To confirm purity, fractions were run on a 13% sodium dodecyl sulfate-polyacrylamide electrophoresis (SDS-PAGE) gel and visualized with Coomassie Brilliant Blue. Selected fractions were further concentrated and flash frozen in liquid nitrogen for further use.

### Protein expression and purification of Eag-TMD complexes

Eag-TMD complexes and their point variants were expressed in BL21(DE3)-Gold cells using LB culture medium supplemented with 100 μg/mL ampicillin. Eag-TMD complexes and point variants were lysed and harvested as described above. For Eag-TMD complexes, the Eag-TMD lysis buffer consisted of 50 mM Tris, 500 mM NaCl and 20 mM imidazole. Protein complexes were purified by affinity chromatography using a nickel-NTA agarose gravity column (Goldbio) equilibrated with 1 x column volume of lysis buffer. The bound proteins were washed with 3 x column volumes of lysis buffer before being eluted by 1 x column volume of elution buffer (50 mM Tris pH 8, 500 mM NaCl, 500 mM Imidazole). Following metal-affinity chromatography, Eag-TMD complexes and point variants were further concentrated and purified by SEC using the same protocol as the Eag chaperones. For all Eag-TMD complexes and point variants the gel filtration buffer was 50 mM Tris pH 8, 250 mM NaCl, 1mM BME. Finally, pure fractions as assayed by Coomassie-stained SDS-PAGE gel were concentrated and flash frozen in liquid nitrogen for further crystallization and biophysical experiments.

### Dynamic Light Scattering (DLS)

Dynamic light scattering (DLS) experiments were performed with a Prometheus Panta (NanoTemper) in DLS buffer (50 mM Tris pH 8.0, 250 mM NaCl, 1 mM BME) at 25°C and a protein concentration of 0.5 mg/ml. Each measurement lasted for 15 s and was repeated 10 times in triplicate thus providing accumulation of 30 measurements. The calculations were performed with the Panta Analysis software using the intensity distributions for all subsequent evaluations.

### Nano-Differential Scanning Fluorimetry (nanoDSF)

Thermal unfolding experiments of purified wild-type and variant Eag chaperones and Eag- TMD complexes were performed in a Prometheus NT.48 instrument (NanoTemper Technologies Inc.). Protein samples were diluted in gel filtration buffer (50 mM Tris pH 8, 250 mM NaCl, 1mM BME) to 0.5 mg/mL and loaded in 10 μL nanoDSF Grade Standard capillaries in triplicate. Samples were heated from 20 °C to 95 °C with a temperature ramp of 1.0 °C/min. Florescence intensity was monitored at 330nm and 350nm after excitation of tryptophan at 280nm. Melting temperatures (T_m_) were calculated from the first derivative of the 350/330nm emission ratio. Data was analyzed with the PR. ThermControl software (NanoTemper Technologies Inc.).

### Differential Scanning Calorimetry (DSC)

Purified wild-type Eag chaperones and Eag-TMD complexes were dialyzed overnight into buffer containing 50 mM PBS (pH 7.5), 250 mM NaCl, and 0.1 mM TCEP. For DSC measurements, an equal volume of 0.5 mg/mL protein solution in DSC buffer as well as a reference solution containing DSC buffer without the addition of protein were prepared. Both the protein and reference solutions were degassed for 10 minutes under vacuum to ensure removal of air bubbles prior to loading of DSC. Experiments were performed using a Nano DSC (TA Instruments, New Castle DE) with a 0.3 mL capillary cell volume. Samples were equilibrated at 20°C for 600 s and heated to 95°C with a scan rate of 1°C per minute at a constant pressure of 3 atm. A background scan was performed by loading both sample and reference cells with reference solution prior to the sample scan. Data were analyzed using NanoAnalyze (TA Instruments, New Castle DE). The sample background was subtracted, and a sigmoidal baseline was applied to the data. The data was fit to a two-state scaled model for the transition.

### Steady-state tryptophan fluorescence spectral analysis

For chemical denaturation experiments using the Prometheus NT.48 (fixed 330 and 350 nm), purified proteins were denatured in increasing concentrations of either urea of guanidine hydrochloride (0.08M to 7.6M urea, Δ dilution = 0.3 per capillary; 0.6M to 5.6M guanidine hydrochloride, Δ dilution = 0.24 per capillary). For each experiment, capillaries containing 24 different concentrations of denaturant and 0.5 mg/mL of protein were loaded in triplicate into a Prometheus NT.48 instrument (NanoTemper Technologies Inc.). Data was analyzed with the PR. ChemControl software (NanoTemper Technolgies Inc.)

For quantitative spectral analysis of wild-type Eags and both Q58A and L66A variants, intrinsic protein fluorescence was measured in the presence of increasing concentrations of guanidine hydrochloride (GdmHCl) (1.50M, 2.0M, 3.0M, 3.5M, 4.0M, 4.5M, 5.0M, 5.5M, 6.0M). Concentrated protein was dialyzed in 1 L of buffer (25 mM Tris pH 7.0) for 1.5 hours to ensure adequate buffer exchange. The protein was then diluted to the appropriate stock concentration. Steady-state fluorescence spectra were measured on a Fluorolog-3 Horiba Jobin Yvon spectrofluorometer (Edison, NJ) using a 10 × 2mm^2^ quartz cuvette to hold the sample. The samples were excited at 280 nm, the excitation and emission slits were set to a 2 nm bandpass. A series of 500 μL samples with a final concentration of 1μM protein dimer (2 μM monomer), were prepared at various GdmCl concentrations. All samples were prepared in triplicate. The samples were equilibrated at room temperature for 1 hour prior to the scan. The final denaturant concentrations were determined by refractive index using C10 Abbe Refractometer (VEE GEE Scientific) post data acquisition. The data were analyzed with Sigma Plot (Point Richmond, CA) software.

### Calculation of free energy of Eag unfolding and TMD binding from steady-state tryptophan fluorescence

If it is assumed that the chaperones SciW and EagT6 unfold through a two-state mechanism:

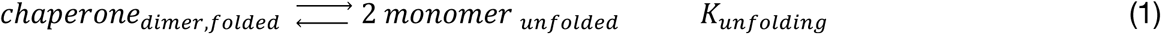

The fraction of tryptophan residues in the unfolded state “α” can be related to the measured *λ_max_* through the equation^45^:

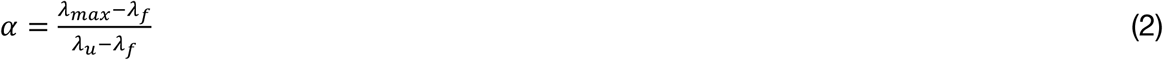

The denaturant concentration dependencies of *λ_f_* and *λ_u_* are given in Figure S3. Allowing us to define the equilibrium constant:

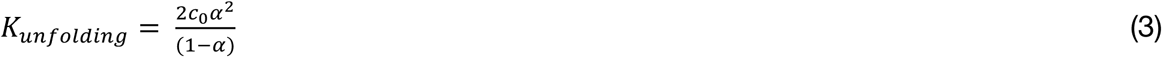

Where *c*_0_ is the the total concentration of Eag monomer in solution, *λ*_f_ is the emission maximum wavelength of the folded protein and *λ*_u_ is maximum wavelength of the unfolded protein.

Next, Kunfolding is related to the change in free energy by:

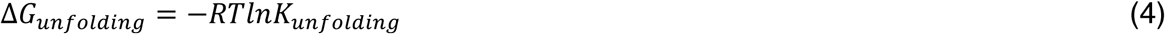

Which allows the determination of the standard change in unfolding free energy in buffer 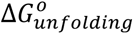, assuming a linear free energy dependence upon denaturation concentration^46^:

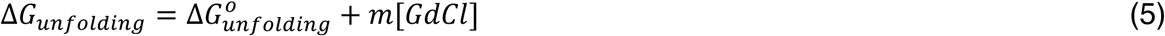

Based upon this, the denaturation concentration dependence of α (fraction unfolding) is given by:

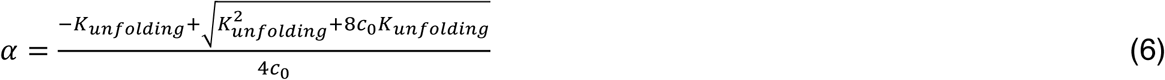

In which 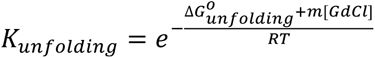.

If the TMD is assumed to selectively bind to the folded form of the Eag with an association constant *K_TMD_*, the effect of TMD binding upon the standard unfolding free energy for the chaperone can be estimated using the relationship^48,49^:

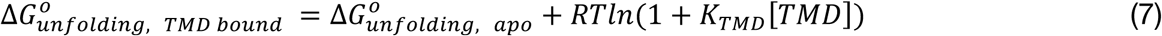

Where 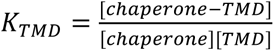, if the TMD binds the chaperone tightly, the majority of the chaperone is bound to the TMD and [*chaperone* − *TMD*] ≈ 0.5 *c*, 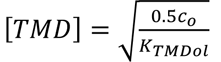 and *K_TMD_*[*TMD*] is significantly larger than one. We can then simplify Equation 7 to:

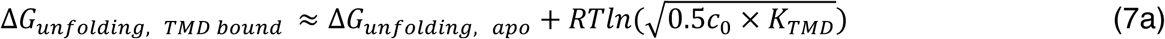

Rearranging this equation we obtain:

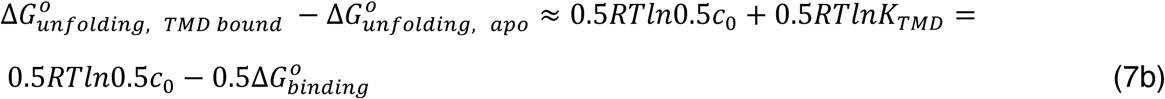

### Stopped-flow kinetics

Unfolding kinetics were performed using Applied photophysics SX-20 (Surrey, UK) stopped-flow fluorescence instrument (dead time <2 ms). To ensure the appropriate dilution of denaturant, asymmetric mixing was set up using 2.5 mL and 250 uL drive syringes purchased from Delta photonics (Ottawa, CA). Purified wild-type and variant Eag chaperones and complexes were first diluted to a concentration of 10 μM in gel filtration buffer (50 mM Tris, pH 8, 250 mM NaCl and 1 mM BME). Purified protein was diluted in a 1/10, v/v, mixing ratio with guanidine hydrochloride (GdmHCl) in 50 mM Tris, 250 mM NaCl, 1 mM BME of the desired concentration. The excitation wavelength was set to 280 nm, and the emission was monitored using a 330 nm Bandpass filter (FWHM 10nm). All data was collected at 20 °C. At least 12 unfolding traces were collected at each desired concentration of GdmCl and averaged. The final denaturant concentrations were determined by refractive index using C10 Abbe Refractometer (VEE GEE Scientific) post data acquisition. All data was analyzed using Sigma Plot 12 software (Point Richmond, CA).

### Calculation of unfolding rate constants and psi-value analysis from stopped-flow kinetics

The EagT6 and SciW relaxation profiles can be described as a mono-exponential decay:

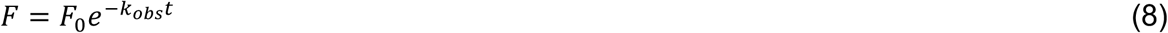

Where *F* is the measured fluorescence, *F*_0_ is the fluorescence at time zero and *k_obs_* is the measured rate constant; at high denaturant concentrations *k_obs_* ≈ *k_unfolding_*, *k_unfolding_* being the apparent unfolding rate constant. Using a chevron-type analysis, it can be assumed:

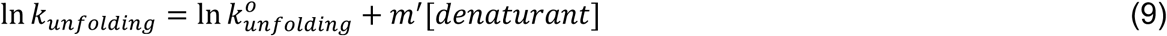

Where *k_unfolding_* is the unfolding rate constant measured at a given concentration of denaturant, ln *k°_unfolding_* is the unfolding rate constant in pure buffer, and *m*′ is the “kinetic m-value”. We have measured time-dependent unfolding traces of all Eag proteins (WT and variants), in the apo and TMD-bound states and have plotted ln(*k_obs_* ≈ *k_unfolding_*) as a function of denaturant concentration in Figure 5; analyzing this date via Eq-7 yields the kinetic parameters ln *k^o^_unfolding_* and *m*′ as shown in Table 1.

For “φ-value analysis”, if the free energy of the unfolding pathway’s rate-limiting transition state is represented by 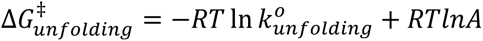 in which *A* is the pre-exponential factor defined by transition-state theory and the φ-value for each mutant is defined as:

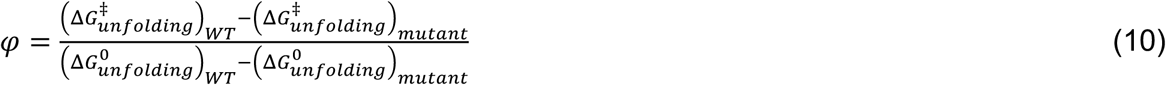

### *P. aeruginosa* strains and growth conditions

*P. aeruginosa* strains were derived from the sequenced strain PAO1. Cultures were grown in lysogeny broth (LB) medium (10 g/L tryptone, 5 g/L yeast extract, 10 g/L NaCl) shaking at 37°C. Solid media containing 1.5% or 3% (w/v) agar, as indicated. *E. coli* strains XL1 blue and SM10 were used for plasmid maintenance and conjugative transfer, respectively. *E. coli* strains were grown in LB at 37°C in a shaking incubator. Cultures were supplemented with 15 mg/mL gentamicin (*E. coli*), 30 mg/mL gentamicin (*P. aeruginosa*), and 25 mg/mL irgasan (*P. aeruginosa*).

### DNA manipulation, plasmid construction, and strain generation for bacterial competition and secretion assays

All primers were synthesized by Integrated DNA Technologies (IDT). Phusion polymerase, restriction enzymes, and T4 DNA ligase were obtained from New England Biolabs (NEB). Sanger sequencing was performed by the Centre for Applied Genomics, The Hospital for Sick Children in Toronto, Ontario. Chromosomal mutations in *P. aeruginosa* were generated by double allelic exchange as previously described^64^. Approximately 500 bp flanks upstream and downstream of the region to be mutated were amplified by PCR and spliced together by overlap-extension PCR. Primers were designed such that the desired point mutation was contained within the overlap. The resulting amplicon was ligated into the pEXG2 allelic exchange vector, transformed into *E. coli* SM10, and introduced to *P. aeruginosa* by conjugative transfer. Merodiploids were selected on LB agar containing 30 mg/mL gentamicin and 25 mg/mL irgasan and streaked on LB agar lacking NaCl containing 5% (w/v) sucrose for *sacB* counterselection. Strains that grow on sucrose but are gentamicin sensitive were screened by Sanger sequencing the PCR amplicon of the region of interest to confirm the presence of point mutations.

### Bacterial competition assays

Assays were performed in co-culture as previously described^65^. Recipient strains harboured a constitutely expressed lacZ gene at a neutral site in the chromosome to distinguish them from donor strains upon plating on LB containing 40 mg/mL 5-bromo-4- chloro3-indolyl-β-D-galactopyranoside (X-gal). Overnight cultures of indicated donor and recipient strains were diluted to an OD600 of 1.0 and combined in an initial donor/recipient ratio of 5:1(v/v), and the initial mixture was serially diluted to enumerate colony forming units (CFU) by plating on media containing X-gal. 10 *μ*L of this mixture was spotted onto a nitrocellulose membrane on LB containing 3% (w/v) agar and cultured at 37°C for 20 hours. Co-cultures were resuspended in 1 mL LB and serially diluted to enumerate CFU. Data are presented as the fold change in the ratio of predator to prey at the end of the experiment relative to the beginning.

### Protein crystallization

Purified SciW-Rhs1(1-59) L66A was screened for crystallization utilizing commercially available screens (Nextal) and a crystal Gryphon robot (Art Robbins Instruments). Crystals of SciW-Rhs1 (1-59) L66A grew in gel filtration buffer and 0.1 M MES pH 6, 0.2 M MgCl_2_, and 20% (v/v) PEG 6000 20 mg/mL protein in a 1:1 mixture at 4 °C. Crystals were further optimized by micro-seeding using a SeedBead kit (Hampton Research).

### Data Collection and Refinement

A data set for SciW-Rhs1(1-59) L66A was collected at the Canadian Light Source beamline CMCF-BM (08B1). Protein crystals were first cryo-protected in 30% PEG 6000 before flash freezing directly in liquid nitrogen. Data was processed using XDS^66^ and CCP4^67^. Initial phases were obtained from Phenix^68^using SciW-Rhs1(1-59) structure (PDBid 6XRR) as a model for molecular replacement. The structure was further built using Coot^69^ and refined using Phenix, and TLS refinement^70^. Molecular graphics were generated using ChimeraX and GraphPad Prism version 10.0.2.

## DATA AVAILABILITY

The X-ray structure and diffraction data reported in this paper have been deposited in the Protein Data Bank under the accession codes 9MVG.

## SUPPORTING INFORMATION

This article contains supporting information.

## ACKNOWLEGEMENTS

We would like to thank the laboratory of Jörg Stetefeld at the University of Manitoba for DLS instrument access. We would also like to thank beamline CMCF-BM at the Canadian Light Source, which is supported by the Canada Foundation for Innovation (CFI), the Natural Sciences and Engineering Research Council (NSERC), the National Research Council (NRC), the Canadian Institutes of Health Research (CIHR), the Government of Saskatchewan.

## AUTHOR CONTRIBUTIONS

M.V.S and G.P. conceived the study. M.V.S expressed, purified, and crystalized proteins. A.G expressed and purified proteins. M.V.S and I.A performed biophysical experiments, and M.V.S, I.A., and M.K. analyzed all biophysical experiments. M.V.S and G.P solved and analyzed the crystal structure. J.C. performed bacterial competition assays and S.A. and J.C. constructed the Pseudomonas strains used in this study. M.V.S, I.A, M.K, J.C.W, and G.P wrote the paper. All authors provided feedback on the manuscript.

## FUNDING AND ADDITIONAL INFORMATION

This work was supported by a Canadian Institutes of Health Research (CIHR) project grant PJT-180450 to G.P, a Research Manitoba New Investigator Operating grant to G.P., and a Canadian Foundation for Innovation (CFI) grant 37841 to G.P. This work is also supported by Canadian Institute of Health Research (CIHR) grant PJT-175011 to J.C.W and a National Sciences and Engineering Research Council of Canada (NSERC) Discovery Grant RGPIN-2024-06410 to M.K. M.V.S is supported through a Research Manitoba Graduate Studentship and University of Manitoba Graduate Fellowship (UMGF). I.A. is also supported through a University of Manitoba Graduate Fellowship (UMGF).

