## Supplemental_data for "Biophysical characterization of Eag chaperones suggests the mechanism of effector transmembrane domain release"

Running title: An Eag conformational change releases transmembrane domains

**Key words:**
Type VI secretion system, Eag, chaperone, membrane protein, stopped-flow, protein folding, X-ray crystallography

*To whom correspondence should be addressed: G.P.

Telephone: (+1) 204-474 -6543

Tables S1 to S5
Figures S1 to S10
Movie S1 (Description in this PDF)

**Table S1: NanoDSF thermal and chemical denaturation values and fit paraments of wild-type Eag and Eag:TMD complexes.**

| Protein | Tm | | Urea | | GdmCl | |
|---|---|---|---|---|---|---|
| | °C | $R^2$ | $C_{50}$ [M] | $R^2$ | $C_{50}$ [M] | $R^2$ |
| **SciW** | 56.7 | 0.917 | 3.41 | 0.991 | 1.45 | 0.995 |
| **SciW-TMD** | 86.6 | 0.988 | N/A | N/A | 3.45 | 0.984 |
| **EagT6** | 53.8 | 0.947 | 3.34 | 0.997 | 1.03 | 0.993 |
| **EagT6-TMD** | 68.7 | 0.970 | 5.80 | 0.994 | 1.95 | 0.990 |

The melting temperature (Tm) was fit to a gaussian distribution and coefficients of determination ($R^2$) provided. The urea and guanidine hydrochloride (GdmCl) denaturation’s were fit to a variable slope model and subsequent $C_{50}$ [M] values provided. The coeffects of determination ($R^2$) are > 0.98 suggesting the unfolding of Eag and Eag-TMD complexes are well-described by a variable slope model.

**Table S2: NanoDSF thermal and chemical denaturation values and fitting parameters for Eag and Eag:TMD point variants.**

| Mutant | Protein | GdmCl | |
|---|---|---|---|
| | | $C_{50}$ [M] | $R^2$ |
| I24/5 F | SciW I25F | 1.05 | 0.995 |
| | SciW-TMD I25F | 2.80 | 0.992 |
| | EagT6 I24F | 0.88 | 0.995 |
| | EagT6-TMD I24F | 0.99 | 0.992 |
| S41A | SciW S41A | 1.44 | 0.974 |
| | SciW-TMD S41A | 3.20 | 0.989 |
| | EagT6 S41A | 0.84 | 0.993 |
| | EagT6-TMD S41A | 1.38 | 0.992 |
| Q58A | SciW Q58A | 1.24 | 0.993 |
| | SciW-TMDQ58A | 2.72 | 0.995 |
| | EagT6 Q58A | 1.00 | 0.997 |
| | EagT6-TMD Q58A | 1.37 | 0.994 |
| L66A | SciW L66A | 1.21 | 0.996 |
| | SciW-TMD L66A | 3.11 | 0.992 |
| | EagT6 L66A | 0.72 | 0.989 |
| | EagT6-TMD L66A | 0.64 | 0.978 |
| Q102A | EagT6 Q102A | 0.96 | 0.995 |
| | EagT6-TMD Q102A | 1.00 | 0.994 |

Thermal denaturation curves were fit to a gaussian distribution and coefficients of determination ($R^2$) values are provided. Chemical denaturation values. All proteins were subjected increasing concentrations of GdmCl and the denaturation profiles were fit to a variable slope model with $C_{50}$ [M] coefficients of determination ($R^2$) values provided.

**Table S3: Strains used in this study.**

| Organism | Genotype | Description | References |
|---|---|---|---|
| *Pseudomonas aeruginosa* PAO1 | Wild type | | [1] |
| | ΔPA4856 | *retS* deletion strain | [2] |
| | ΔPA4856 ΔPA0093 ΔPA0092 attB::*lacZ* | *retS, tse6/tsi6* deletion strain | [3] |
| | ΔPA4856 ΔPA0094 | *retS, eagT6* deletion strain | [4] |
| | ΔPA4856 PA0094_I24F | *retS* deletion, *eagT6* I24F mutation | This study |
| | ΔPA4856 PA0094_S41A | *retS* deletion, *eagT6* S41A mutation | This study |
| | ΔPA4856 PA0094_Q58A | *retS* deletion, *eagT6* Q58A mutation | This study |
| | ΔPA4856 PA0094_L66A | *retS* deletion, *eagT6* L66A mutation | This study |
| | ΔPA4856 PA0094_Q102A | *retS* deletion, *eagT6* Q102A mutation | This study |
| *E. coli* XL-1 Blue | *recA1 endA1 gyrA96 thi-1 hsdR17 supE44 relA1 lac* [F´ *proAB lacI*$^{q}$ ZΔ*M15* Tn*10* (Tet$^{R}$)] | Cloning strain | Novagen |
| *E. coli* SM10 λpir | *thi thr leu tonA lac Y supE recA*::RP4-2-Tc::Mu | Conjugation strain | BioMedal LifeScience |
| *E. coli* BL21 (DE3) CodonPlus | F$^{-}$ ompT gal dcm lon hsdS$_{B}$(r$_{B}^{-}$ m$_{B}^{-}$) λ(DE3) pLysS(cm$^{R}$) | Protein expression strain | Novagen |

**Table S4: Plasmids used in this study**

| Plasmid | Relevant features | Reference |
|---|---|---|
| pETDuet-1:: SL1344_0286_1-59-His$_6$:: SL1344_0285-VSV-G | Co-expression vector for C-terminal His$_6$ tagged Rhs1 TMD and C-terminal VSV-G tagged SciW | 5 |
| pETDuet-1::PA0093_1-61-His$_6$::PA0094-VSV-G | Co-expression vector for C-terminal His$_6$ tagged Tse6 TMD1 and C-terminal VSV-G tagged EagT6 | 5 |
| pRSETA::SL1344_0285 | Expression vector for SciW from *Salmonella* Typhimurium (for crystallization) | 5 |
| pET29b::SL1344_0285-VSVG | Expression vector for SciW | This study |
| pET28a::PA0094 | Expression vector for EagT6 | |
| pET29b::PA0094-VSV-G | Expression vector for C-terminal VSV-G tagged EagT6 from *P. aeruginosa* | 5 |
| pETDuet-1:: SL1344_0286_1-59-His$_6$:: SL1344_0285-VSV-G I24F | *sciW*_TMD I24F mutation construct | This study |
| SL1344_0286_1-59-His$_6$:: SL1344_0285-VSV-G S41A | *sciW*_TMD S41A mutation construct | This study |
| SL1344_0286_1-59-His$_6$:: SL1344_0285-VSV-G Q58A | *sciW*_TMD Q58A mutation construct | This study |
| SL1344_0286_1-59-His$_6$:: SL1344_0285-VSV-G L66A | *sciW*_TMD L66A mutation construct | This study |
| pETDuet-1::PAA0093_1-61-His6::PA0094-VSV G I24F | *eagT6*_TMD I24F mutation construct | This study |
| pETDuet-1::PAA0093_1-61-His6::PA0094-VSV G S41A | *eagT6*_TMD S41A mutation construct | This study |
| pETDuet-1::PAA0093_1-61-His6::PA0094-VSV G L66A | *eagT6*_TMD L66A mutation construct | This study |
| pETDuet-1::PAA0093_1-61-His6::PA0094-VSV G Q58A | *eagT6*_TMD Q58A mutation construct | This study |
| pETDuet-1::PAA0093_1-61-His6::PA0094-VSV G Q102A | *eagT6*_TMD Q102A mutation construct | This study |
| pET29b::SL1344_0285-VSVG I24F | *sciW* I24F mutation construct | This study |
| pET29b::SL1344_0285-VSVG S41A | *sciW* S41A mutation construct | This study |
| pET29b::SL1344_0285-VSVG Q58A | *sciW* Q58A mutation construct | This study |
| pET29b::SL1344_0285-VSVG L66A | *sciW* L66A mutation construct | This study |
| pET29b::PA0094-VSV-G I24F | *eagT6* I24F mutation construct | This study |
| pET29b::PA0094-VSV-G S41A | *eagT6* S41A mutation construct | This study |
| pET29b::PA0094-VSV-G Q58A | *eagT6* Q58A mutation construct | This study |
| pET29b::PA0094-VSV-G L66A | *eagT6* L66A mutation construct | This study |
| pET29b::PA0094-VSV-G Q102A | *eagT6* Q102A mutation construct | This study |
| pEXG2 | Allelic replacement vector containing *sacB,* Gm$^R$ | 6 |
| pEXG2::PA0094_I24F | *eagT6* I24F mutation construct | This study |
| pEXG2::PA0094_S41A | *eagT6* S41A mutation construct | This study |
| pEXG2::PA0094_Q58A | *eagT6* Q58A mutation construct | This study |
| pEXG2::PA0094_L66A | *eagT6* L66A mutation construct | This study |
| pEXG2::PA0094_Q102A | *eagT6* Q102 mutation construct | This study |

### Table S5: Primers for the generation of protein constructs

| Expression Construct | Original Construct | F Primer (5' to 3') | R Primer (3' to 5') | Notes |
|---|---|---|---|---|
| **SciW-TMD I24F** | SciW-TMD | CAGCGTCAATTTTTTTATC CTGG | CGATCCGTAAATGTTTCAG | |
| **SciW-TMD S41A** | SciW-TMD | CCTGAATATTGCCCGCGA TACGC | CTGGGCGATGTTCGTTCA TTG | |
| **SciW-TMD L66A** | SciW-TMD | GAAAAAAAATGCCGGTCA GCAACCGG | ATCAGTGCAATCTGGCGG | |
| **SciW-TMD Q58A** | SciW-TMD | TATTGACCGCTTCATTGCA CTGATGAAAAAAAATATCG | TAGGCGGGCAGGTCTTCA | |
| **EagT6-TMD I24F** | EagT6-TMD | GAGCATCAACTTCTTCAAG CTCCCC | TGGTCCTGCCAGGCATCG | |
| **EagT6-TMD S41A** | EagT6-TMD | TTTCGTCATCGCCCGTGA CGCCAGCCAGG | CTGGCTTCGCGGGCGG | |
| **EagT6-TMD Q58A** | | TGTCGCCCGCGCACTGGA AAACGCC | TAGTCGGCGAACGGCGC | |
| **EagT6-TMD L66A** | EagT6-TMD | CGAGAAGCAAGCGCCCG GCTTCAAG | GCGTTTTCCAGTTGGCGG | |
| **EagT6-TMD Q102A** | EagT6-TMD | GATGCTGCGCTTTGTATTC ATCGAGCGCCGCCCGG | AAGTCGCGGCCCTCGCGC | |

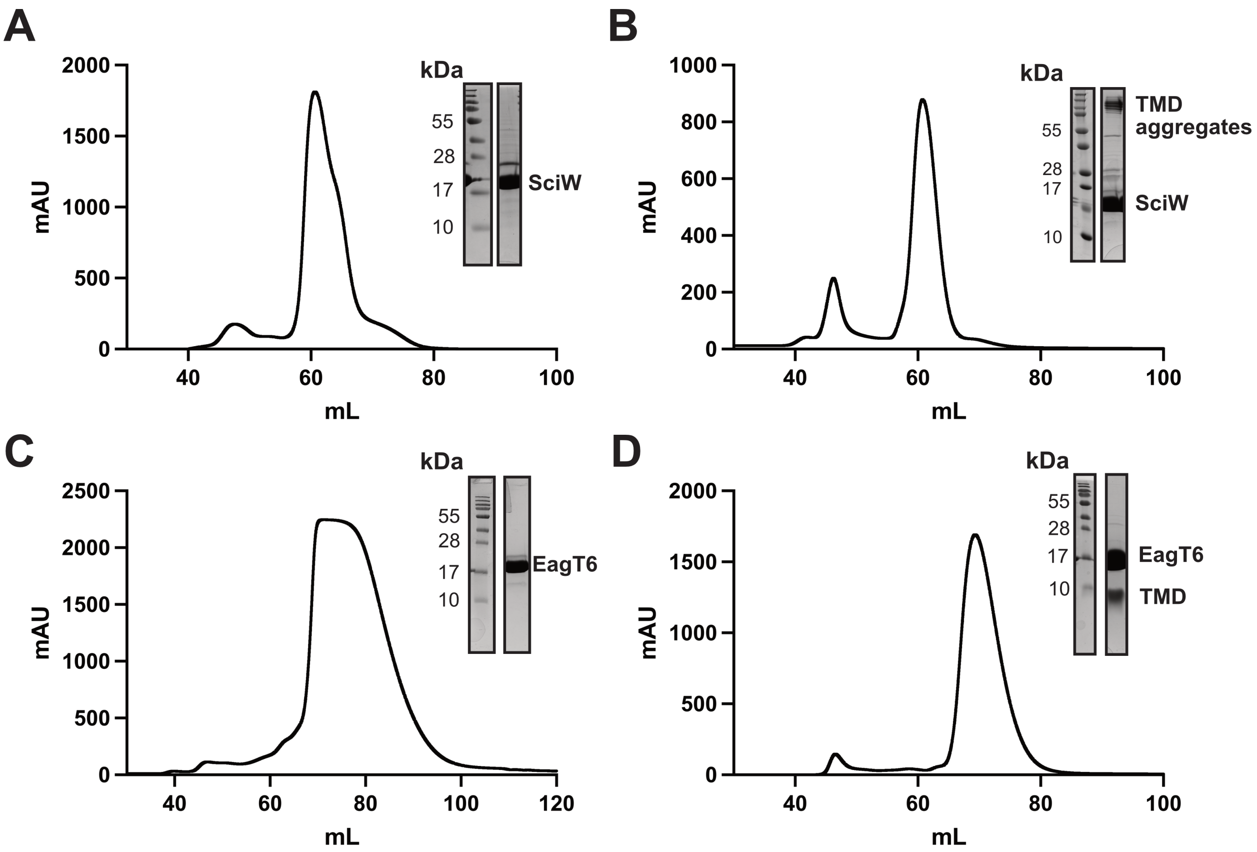


**Figure S1: Purification of Eag and Eag:TMD complexes. (A)** SciW and **(B)** SciW-TMD **(C)** EagT6 and **(D)** EagT6-TMD. All panels show the final purified material after size-exclusion chromatography (SEC) using a HiLoad 16/600 Superdex75 preparatory grade column. The final SEC buffer was 50 mM Tris pH 8.0, 250 mM NaCl, 1 mM BME. To confirm purity, fractions were run on a 13% SDS-PAGE gel and visualized with Coomassie Brilliant Blue. Eag chaperones and TMD bands are marked on each gel.

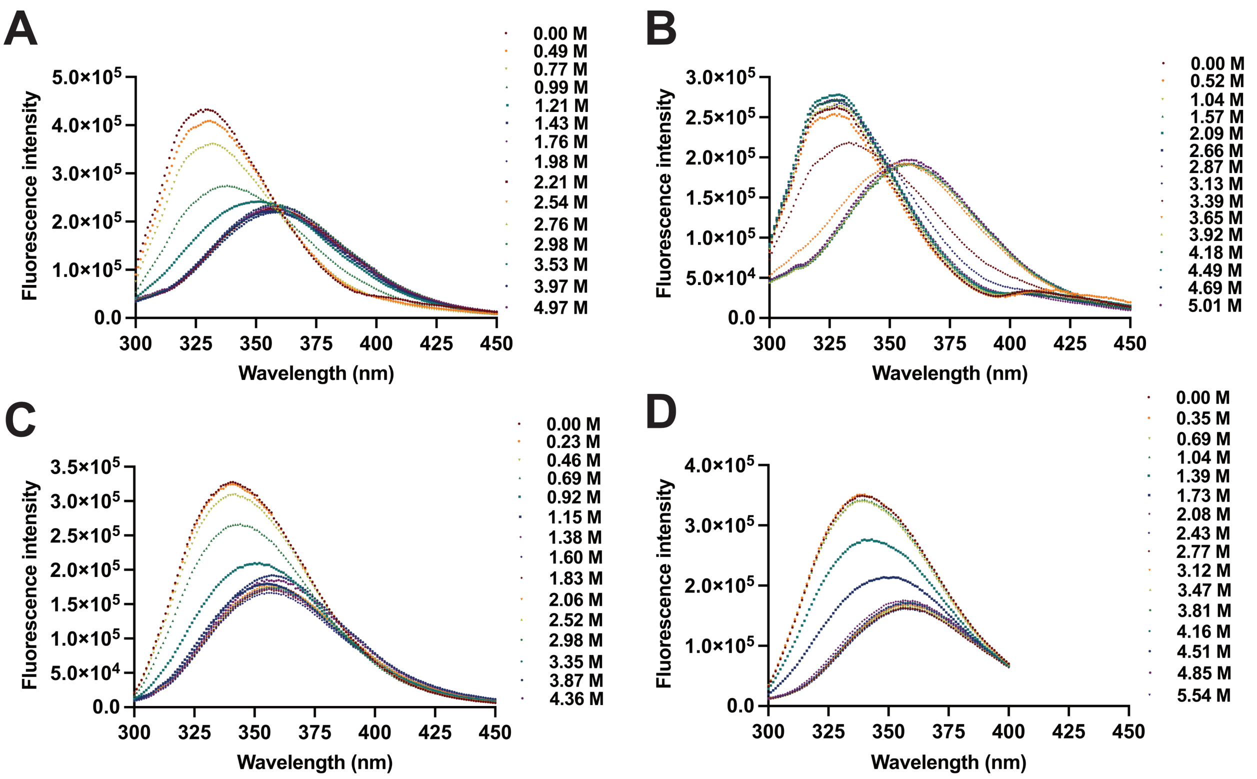


**Figure S2: Steady-state fluorescence spectrum of Eag and Eag-TMD complexes in guanidine hydrochloride.** Steady-state fluorescence spectrum of **(A)** SciW and **(B)** SciW-Rhs1(1-59) and **(C)** EagT6 and **(D)** EagT6-Tse6(1-61) collected in 50 mM Tris (pH 8), 250mM NaCl in increasing concentrations of guanidine hydrochloride. All measurements were performed at room temperature 293 K with a final protein concentration of 10 $\mu M$. The samples were excited at 280 nM with excitation and emission slits set at 2 nM bandpass.

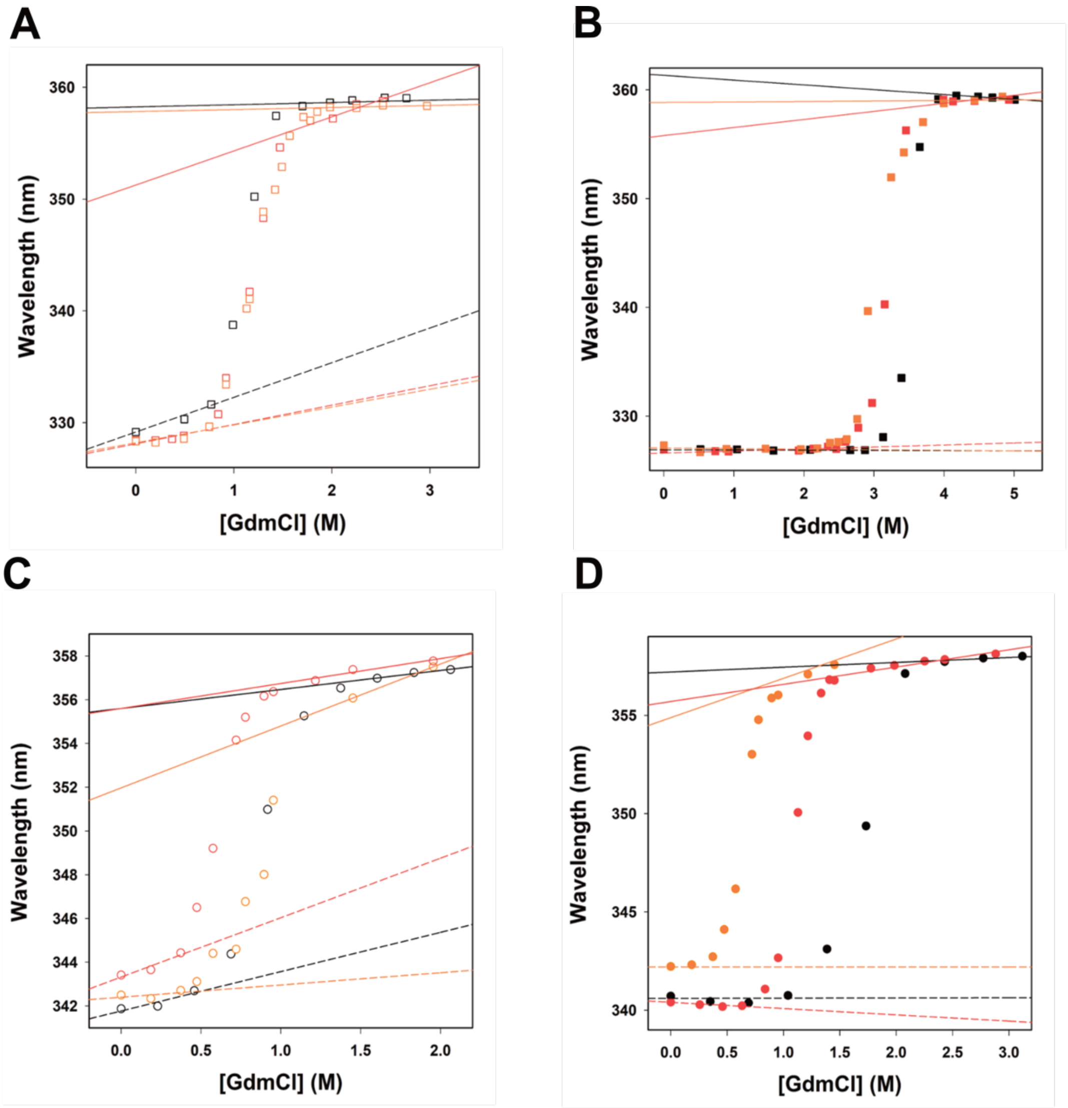


E

| SciW-APO | $\lambda_{folded}$ | $\lambda_{unfolded}$ | SciW-TMD | $\lambda_{folded}$ | $\lambda_{unfolded}$ |
|---|---|---|---|---|---|
| WT | 328.2+1.6[GdmCl] | 358.2+0.2[GdmCl] | WT | 326.9-0.0[GdmCl] | 361.3-0.4[GdmCl] |
| L66A | 328.1+1.7[GdmCl] | 358.8+0.2[GdmCl] | L66A | 327.1-0.1[GdmCl] | 355.8+0.7[GdmCl] |
| Q58A | 328.3-0.5[GdmCl] | 355.0+1.4[GdmCl] | Q58A | 326.6+0.2[GdmCl] | 358.9+0.1[GdmCl] |
| **EagT6-APO** | | | **EagT6-TMD** | | |
| WT | 341.7+1.8[GdmCl] | 355.6+0.9[GdmCl] | WT | 340.6+0.0[GdmCl] | 358.2+0.3[GdmCl] |
| L66A | 343.3+2.7[GdmCl] | 355.6+1.4[GdmCl] | L66A | 340.4-0.3[GdmCl] | 355.7+0.9[GdmCl] |
| Q58A | 342.4+0.6[GdmCl] | 352.0+2.8[GdmCl] | Q58A | 342.2+0.0[GdmCl] | 354.6+2.0[GdmCl] |

**Figure S3: Fraction unfolding fits from steady-state fluorescence data.** Unfolding profile of Eag chaperones, SciW (Squares, panels A and B) and EagT6 (circles, panels C and D) as represented by the lambda max *λmax* of the fluorescence spectra. [Empty shapes = APO, filled shapes = TMD bound, WT is represented in black, Q58A mutant is represented in orange, L66A mutant is represented in red]. It can be clearly seen that these unfolding profiles exhibit three distinct regimes: 1) an initial regime that is linearly dependent on denaturant concentration and represents how the maximum of the folded protein $\lambda f$ changes with denaturant concertation; 2) a nonlinear transition regime indicating the protein unfolding process; 3) a final regime that is also linearly dependent upon denaturation concentration, showing how the maximum of the unfolded protein $\lambda u$ changes with denaturant concentration. (E) Table listing the pre- and post-transition slopes shown in the figure.

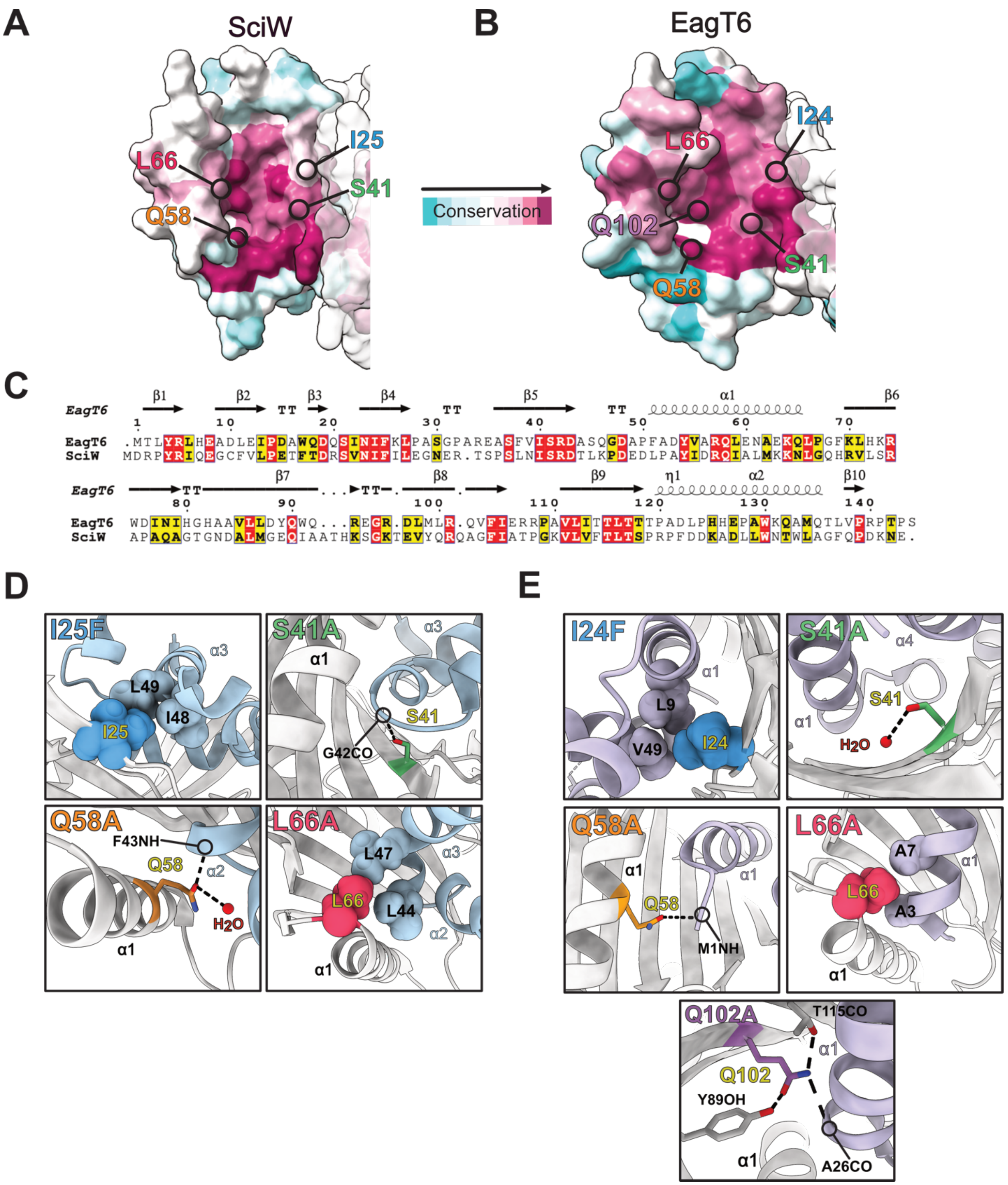


**Figure S4: Conservation and critical residues involved in Eag-TMD binding.** Surface conservation representation of the TMD binding pocket in each chaperone monomer **(A)** SciW and **(B)** EagT6. Conserved residues targeted for point mutation are indicated. **(C)**

Multisequence alignment of SciW and EagT6 as calculated by Consurf (https://consurf.tau.ac.il/). Molecular graphics were drawn using USCF ChimeraX (https://www.rbvi.ucsf.edu/chimerax/). Multisequence alignment was plotted by Espript3 (https://espript.ibcp.fr/). Molecular details of point variant residues involved in binding the TMD in opposing chain shown in Figure 3. **(D)** SciW and **(E)** EagT6 are drawn. Residues are substituted to alanine and colored. I24F (Top Left) creates a hydrophobic surface in the palm of the claw. Hydrophilic residues S41A (Top Right) and Q58A (bottom left) form bifurcated hydrogen bonds with the TMD backbone. L66A (Bottom Right) provides a "knob" for the hole of its cognate TMD.

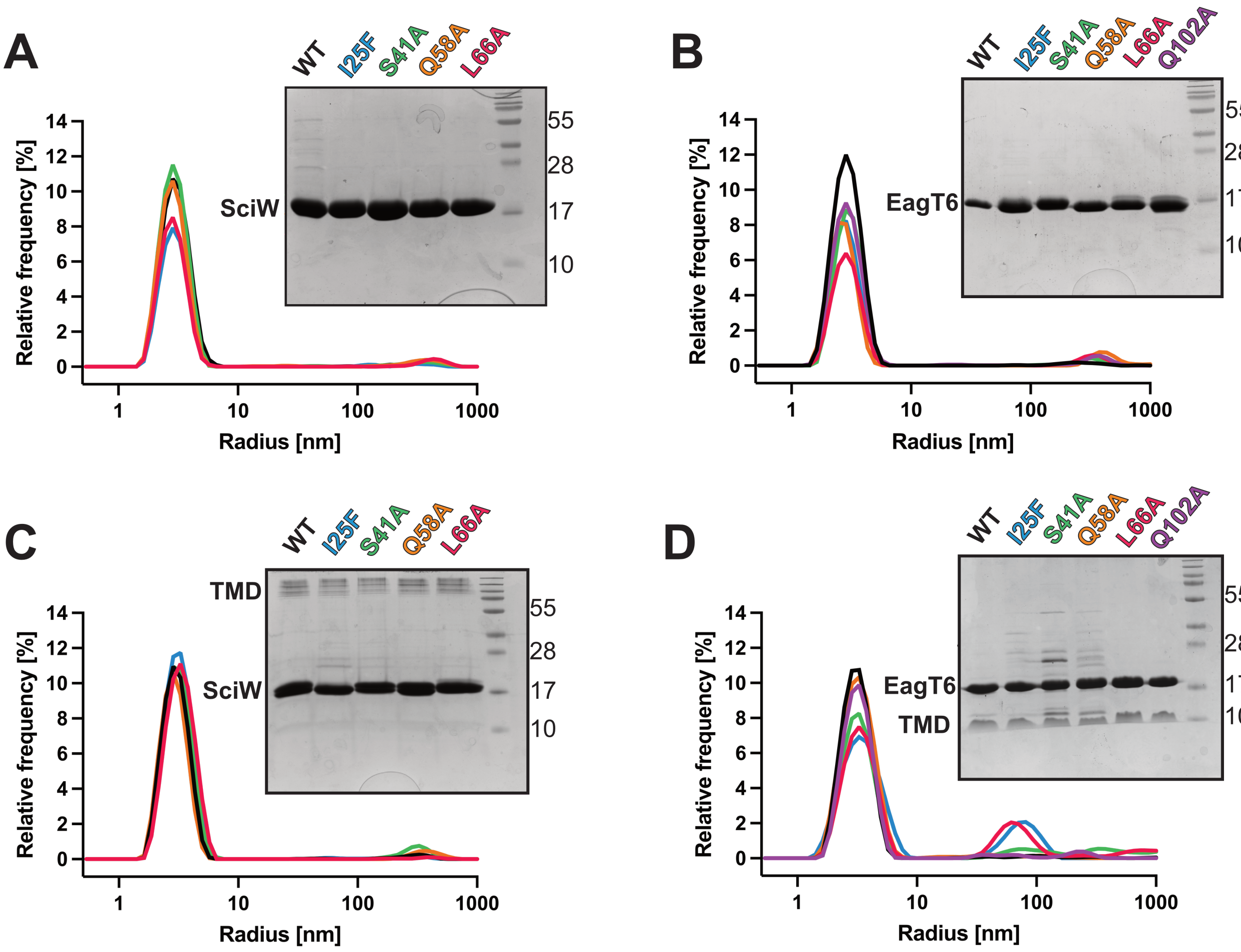


**Figure S5: Eag and Eag:TMD point variants are properly folded and bind effector TMDs.** DLS measured intensity distributions versus radius for **(A)** SciW, **(B)** EagT6, **(C)** SciW-TMD and **(D)** EagT6-TMD. All measurements were collected in 50mM Tris (pH 8), 250mM NaCl and performed at room temperature at a concentration of 0.5 mg/mL in triplicate. Each panel also shows a Coomassie stained SDS-PAGE gel for each purified variant as compared to the wild-type. Molecular weight markers are in kDa.

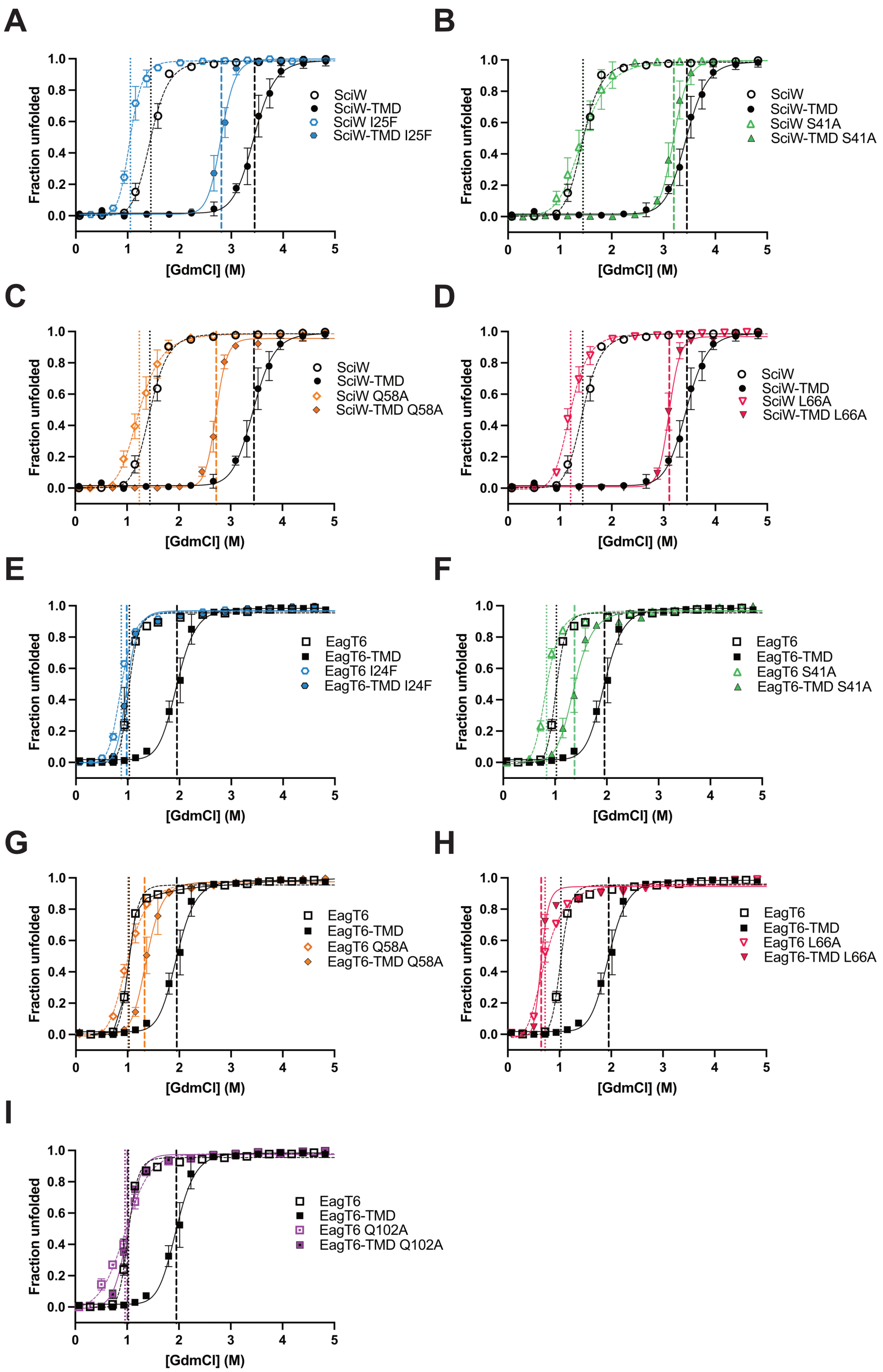
A
B
C
D
E
F
G
H
I
Fraction unfolded
[GdmCl] (M)
SciW
SciW-TMD
SciW I25F
SciW-TMD I25F
SciW S41A
SciW-TMD S41A
SciW Q58A
SciW-TMD Q58A
SciW L66A
SciW-TMD L66A
EagT6
EagT6-TMD
EagT6 I24F
EagT6-TMD I24F
EagT6 S41A
EagT6-TMD S41A
EagT6 Q58A
EagT6-TMD Q58A
EagT6 L66A
EagT6-TMD L66A
EagT6 Q102A
EagT6-TMD Q102A

**Figure S6**: **Chemical denaturation of Eag and Eag:TMD point variants measured by nanoDSF.** Each panel shows an Eag or Eag-TMD point variant relative to the wild-type protein. SciW variants are panels **A-D** and EagT6 variants are panels **E-I**. Apo Eag chaperones (unbound) are represented by unfilled shapes and Eag-TMD complexes by filled shapes. Eag point variants are coloured as follows: I24F (blue), S41A (green), Q58A (orange), L66A (red), wild-type (black). Vertical lines indicate inflection point ($CM_{50}$) and are colored by variant. Error bars represent measurements taken in triplicate.

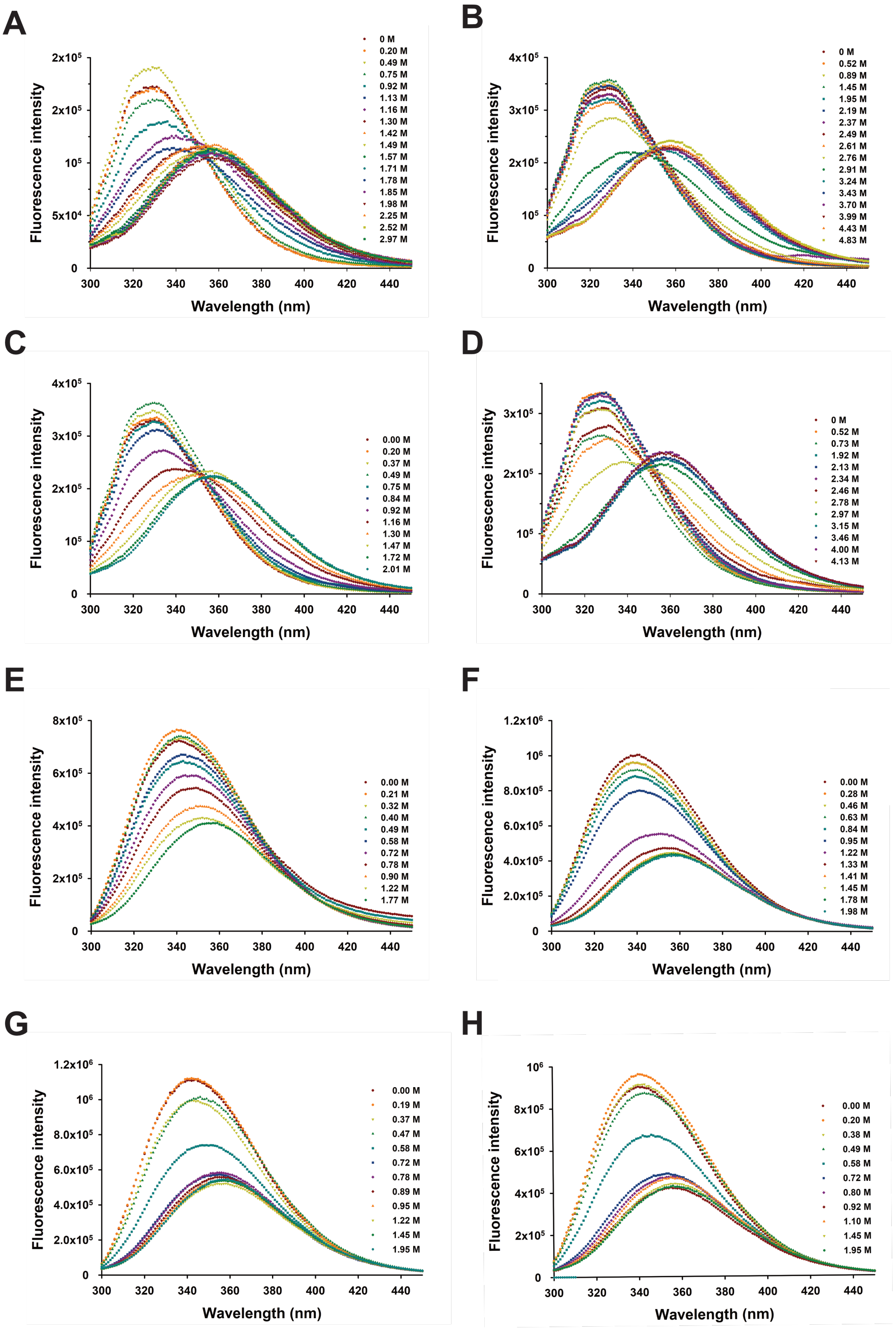
A
Fluorescence intensity
Wavelength (nm)
0 M
0.20 M
0.49 M
0.75 M
0.92 M
1.13 M
1.16 M
1.30 M
1.42 M
1.49 M
1.57 M
1.71 M
1.78 M
1.85 M
1.98 M
2.25 M
2.52 M
2.97 M
B
Fluorescence intensity
Wavelength (nm)
0 M
0.52 M
0.89 M
1.45 M
1.95 M
2.19 M
2.37 M
2.49 M
2.61 M
2.76 M
2.91 M
3.24 M
3.43 M
3.70 M
3.99 M
4.43 M
4.83 M
C
Fluorescence intensity
Wavelength (nm)
0.00 M
0.20 M
0.37 M
0.49 M
0.75 M
0.84 M
0.92 M
1.16 M
1.30 M
1.47 M
1.72 M
2.01 M
D
Fluorescence intensity
Wavelength (nm)
0 M
0.52 M
0.73 M
1.92 M
2.13 M
2.34 M
2.46 M
2.78 M
2.97 M
3.15 M
3.46 M
4.00 M
4.13 M
E
Fluorescence intensity
Wavelength (nm)
0.00 M
0.21 M
0.32 M
0.40 M
0.49 M
0.58 M
0.72 M
0.78 M
0.90 M
1.22 M
1.77 M
F
Fluorescence intensity
Wavelength (nm)
0.00 M
0.28 M
0.46 M
0.63 M
0.84 M
0.95 M
1.22 M
1.33 M
1.41 M
1.45 M
1.78 M
1.98 M
G
Fluorescence intensity
Wavelength (nm)
0.00 M
0.19 M
0.37 M
0.47 M
0.58 M
0.72 M
0.78 M
0.89 M
0.95 M
1.22 M
1.45 M
1.95 M
H
Fluorescence intensity
Wavelength (nm)
0.00 M
0.20 M
0.38 M
0.49 M
0.58 M
0.72 M
0.80 M
0.92 M
1.10 M
1.45 M
1.95 M

**Figure S7: Steady-state fluorescence spectrum of apo Eag and Eag-TMD variants Q58A and L66A.** Steady-state fluorescence spectrum of **(A)** SciW Q58A **(B)** SciW-TMD Q58A **(C)** SciW L66A **(D)** SciW-TMD L66A **(E)** EagT6 Q58A **(F)** EagT6-TMD Q58A **(G)** EagT6 L66A and **(H)** EagT6-TMD L66A collected in 50 mM Tris (pH 8.0), 250mM NaCl in increasing concentrations of guanidine hydrochloride. All measurements were performed at room temperature 293 K with a final protein concentration of 10 µM. The samples were excited at 280 nM with excitation and emission slits set at 2 nM bandpass.

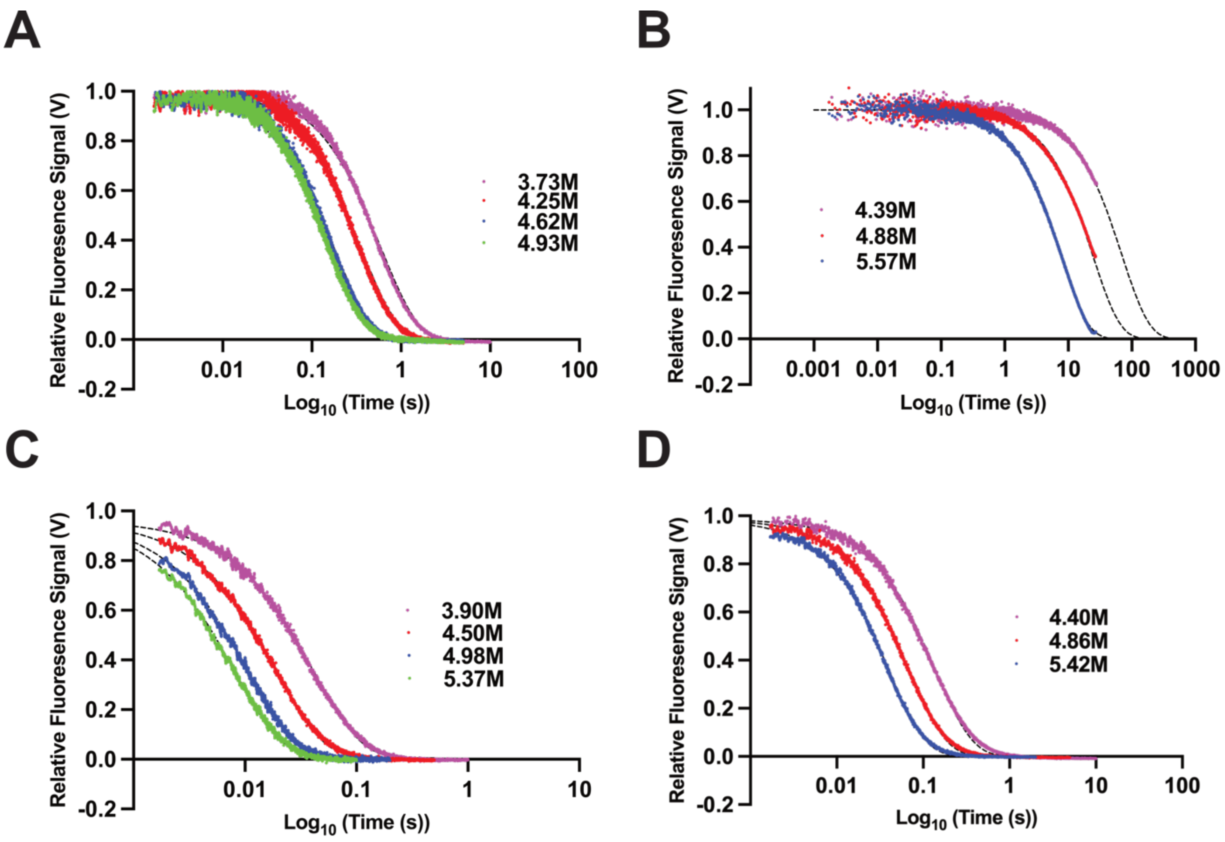


**Figure S8: Fluorescent data for wild-type Eag and Eag:TMD wild-type complex unfolding kinetics** Stopped-flow kinetics traces of **(A)** SciW, **(B)** SciW-TMD **(C)** EagT6, and **(D)** EagT6:TMD in 50 mM Tris (pH 8.0), 250mM NaCl at 20°C. Unfolding traces (fluorescence decrease) are measured in the increasing guanidine concentrations listed. The change in protein fluoresce is monitored at 330 nm and is normalized to 1. The dashed lines represent the best mono-exponential fits to the obtained data.

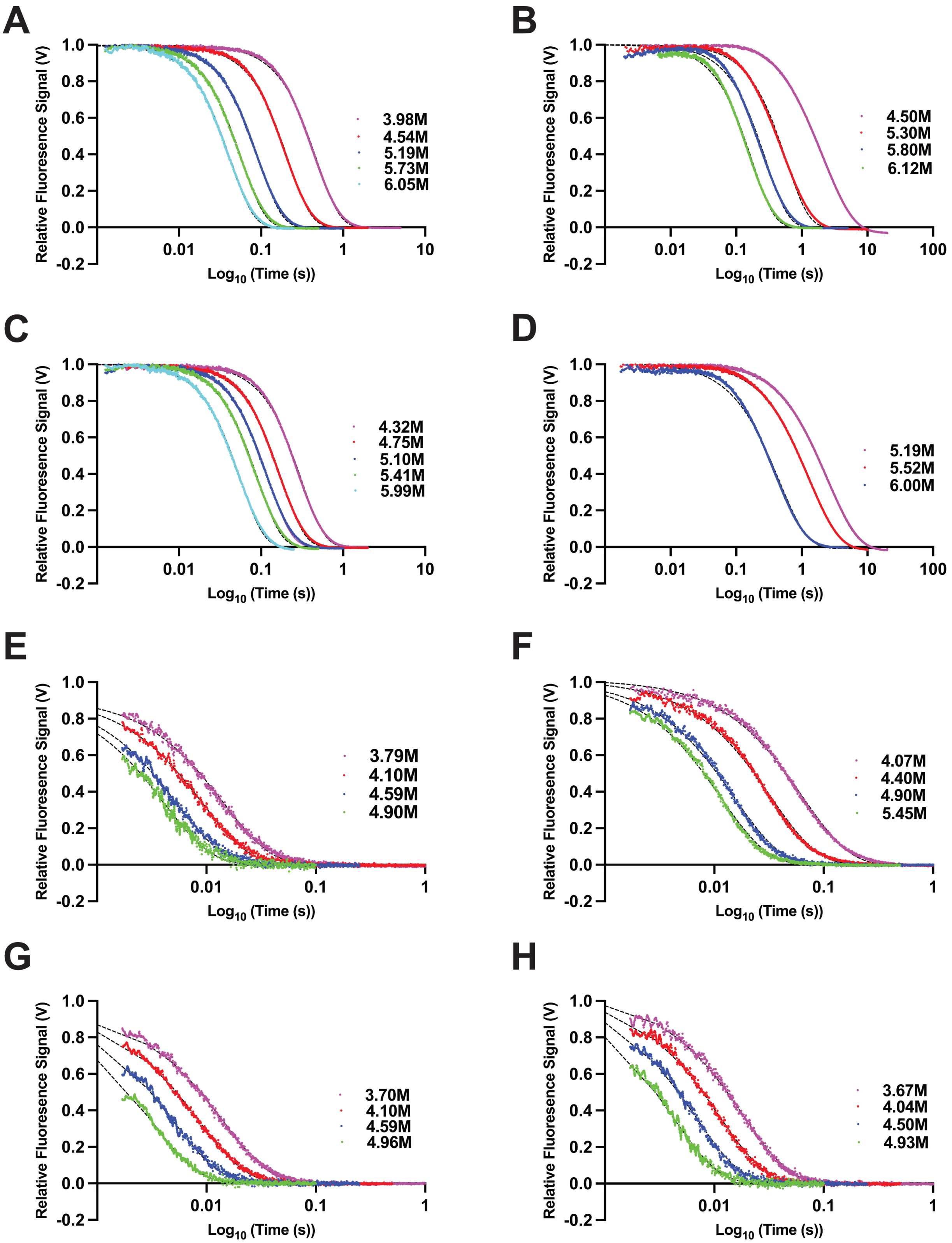
A
Relative Fluoresence Signal (V)
Log10 (Time (s))
3.98M
4.54M
5.19M
5.73M
6.05M
B
Relative Fluoresence Signal (V)
Log10 (Time (s))
4.50M
5.30M
5.80M
6.12M
C
Relative Fluoresence Signal (V)
Log10 (Time (s))
4.32M
4.75M
5.10M
5.41M
5.99M
D
Relative Fluoresence Signal (V)
Log10 (Time (s))
5.19M
5.52M
6.00M
E
Relative Fluoresence Signal (V)
Log10 (Time (s))
3.79M
4.10M
4.59M
4.90M
F
Relative Fluoresence Signal (V)
Log10 (Time (s))
4.07M
4.40M
4.90M
5.45M
G
Relative Fluoresence Signal (V)
Log10 (Time (s))
3.70M
4.10M
4.59M
4.96M
H
Relative Fluoresence Signal (V)
Log10 (Time (s))
3.67M
4.04M
4.50M
4.93M

**Figure S9: Fluorescent data for of Eag and Eag:TMD complex point variant unfolding kinetics** Stopped-flow kinetics traces of **(A)** SciW Q58A **(B)** SciW:TMD Q58A, **(C)** SciW L66A, **(D)** SciW:TMD L66A, **(E)** EagT6 Q58A, **(F)** EagT6:TMD Q58A, **(G)** EagT6 L66A, **(H)** EagT6:TMD L66A in 50 mM Tris (pH 8.0), 250mM NaCl at 20°C. Unfolding traces (fluorescence decrease) are measured in increasing guanidine concentrations listed. The change in protein fluoresce is monitored at 330 nm and is normalized to 1. The dashed lines represent the best mono-exponential fits to the obtained data.

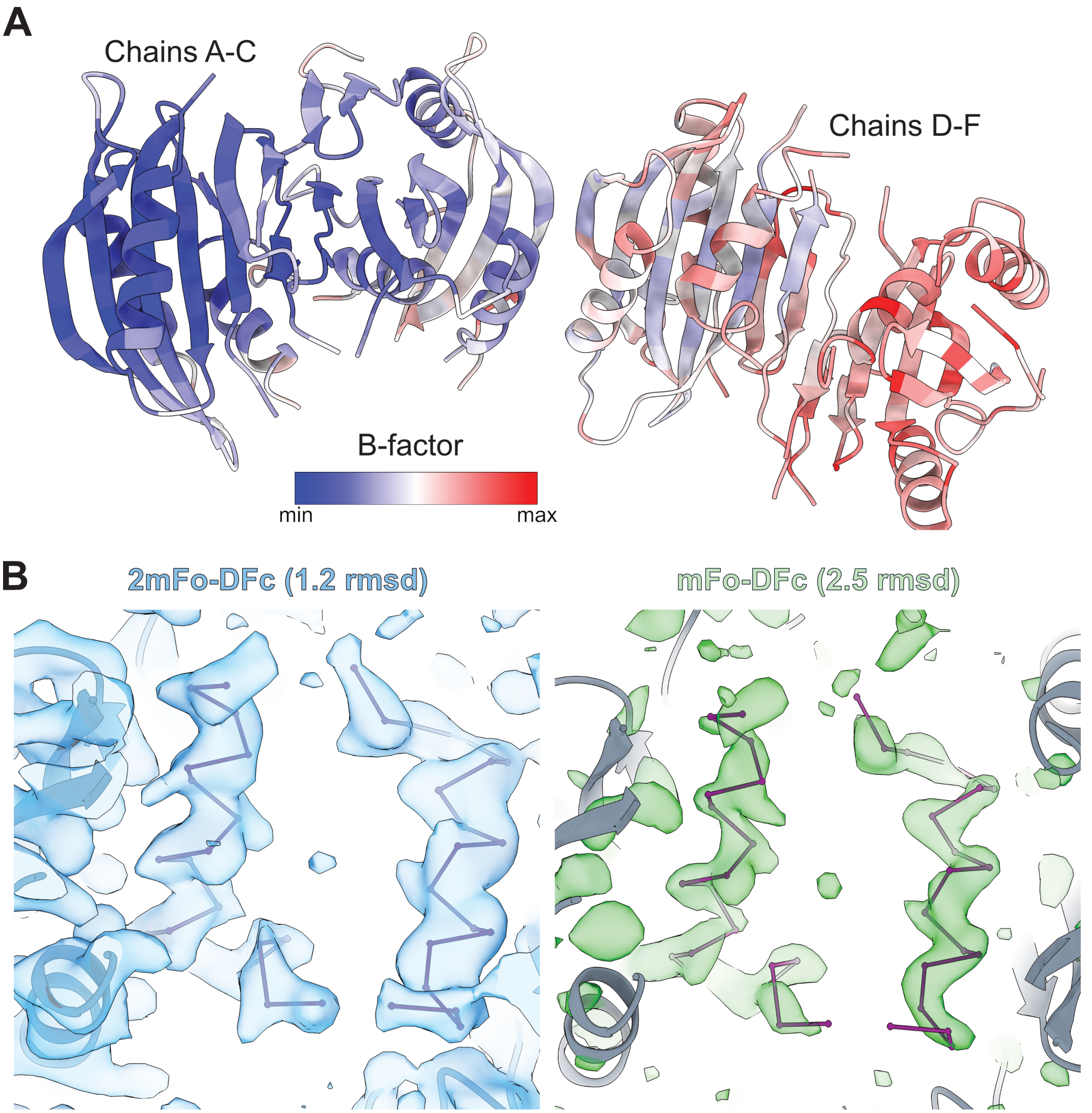


**Figure S10: SciW-TMD L66A structural modeling data. (A)** Asymmetric unit contents colored by B-factor from lowest (blue) to highest (red). **(B)** Electron density maps for the bound TMD in Chains A-C. Right: 2mFo-DFc maps contoured at 1.2 rmsd shown in blue. Left: mFo-DFc difference maps contoured at 2.5 rmsd. Green is positive density. The mFo-DFc map was generated after first omitting chain C (Rhs1 TMD) before refinement.

**Movie S1: Structural morph between wild-type SciW-TMD and SciW-TMD L66A.** Moving showing the conformational change between the wild-type SciW-TMD complex and the partially released TMD in the SciW L66A variant. Movie was made with ChimeraX.
